# Chromatin landscape and epigenetic heterogeneity of acute myeloid leukemia

**DOI:** 10.64898/2026.03.22.711973

**Authors:** Yotaro Ochi, Markus Liew-Littorin, Yasuhito Nannya, Sofia Bengtzen, Benedicte Piauger, Stefan Deneberg, Martin Jädersten, Vladimir Lazarevic, Jörg Cammenga, Anna Robelius, Lovisa Wennström, Emma Ölander, Senji Kasahara, Nobuhiro Hiramoto, Nobuhiro Kanemura, Nobuo Sezaki, Maki Sakurada, Makoto Iwasaki, Junya Kanda, Yasunori Ueda, Satoshi Yoshihara, Tom Erkers, Nona Struyf, Yu Watanabe, Masanori Motomura, Masahiro M Nakagawa, Ryunosuke Saiki, Hidehito Fukushima, Koji Okazaki, Suguru Morimoto, Akinori Yoda, Rurika Okuda, Shintaro Komatsu, Guoxiang Xie, Albin Österroos, Ayana Kon, Lanying Zhao, Yuichi Shiraishi, Takayuki Ishikawa, Satoru Miyano, Shuichi Matsuda, Akifumi Takaori-Kondo, Hiroyuki Aburatani, Hiroshi I Suzuki, Olli Kallioniemi, Gunnar Juliusson, Martin Höglund, Sören Lehmann, Seishi Ogawa

**Affiliations:** Department of Pathology and Tumor Biology, Graduate School of Medicine, Kyoto University, Kyoto, Japan; Institute for the Advanced Study of Human Biology (WPI-ASHBi), Kyoto University, Kyoto, Japan; Center for Hematology and Regenerative Medicine, Department of Medicine Huddinge, Karolinska Institute, Stockholm, Sweden; Division of Hematology, Department of Medicine, Faculty of Medicine and Health, Örebro University, Örebro, Sweden; Division of Hematopoietic Disease Control, Institute of Medical Science, The University of Tokyo, Tokyo, Japan; Department of Hematology, Linköping University Hospital, Linköping, Sweden; Department of Hematology, Oncology and Radiation Physics, Skåne University Hospital, Lund, Sweden; Department of Hematology, Stem Cell Center, Department of Laboratory Medicine, Lund University, Lund, Sweden; Department of Medical Sciences, Hematology, Uppsala University, Uppsala, Sweden; Department of Hematology, Sahlgrenska University Hospital, Gothenburg, Sweden; Department of Hematology, Sundsvall Hospital, Sundsvall, Sweden; Department of Hematology, Gifu Municipal Hospital, Gifu, Japan; Laboratory of Pharmaceutical Health Care and Promotion, Gifu Pharmaceutical University, Gifu, Japan; Department of Hematology, Kobe City Medical Center General Hospital, Kobe, Japan; Department of Hematology & Infectious Disease, Gifu University Hospital, Gifu, Japan; Department of Hematology, Chugoku Central Hospital, Hiroshima, Japan; Department of Hematology, Graduate School of Medicine, Kyoto University, Kyoto, Japan; Department of Hematology/Oncology, Kurashiki Central Hospital, Kurashiki, Japan; Department of Respiratory Medicine and Hematology, Hyogo Medical University, Nishinomiya, Japan; Science for Life Laboratory and Department of Oncology-Pathology, Karolinska Institute, Stockholm, Sweden; Division of Molecular Oncology, Center for Neurological Diseases and Cancer, Nagoya University Graduate School of Medicine, Nagoya, Japan; Department of Nephrology, Nagoya University Graduate School of Medicine, Nagoya, Japan; Division of Hematology and Tumor Biology, Institute of Medical Science, The University of Tokyo, Tokyo, Japan; Division of Stem Cell and Genome Biology, Institute of Medical Science, The University of Tokyo, Tokyo, Japan; Division of Genome Analysis Platform Development, National Cancer Center Research Institute, Tokyo, Japan; Department of Integrated Analytics, M&D Data Science Center, Institute of Integrated Research, Institute of Science Tokyo, Tokyo, Japan; Human Genome Center, Institute of Medical Science, The University of Tokyo, Tokyo, Japan; Department of Orthopaedic Surgery, Graduate School of Medicine, Kyoto University, Kyoto, Japan; Genome Science & Medicine Laboratory, Research Center for Advanced Science and Technology, The University of Tokyo, Tokyo, Japan; Institute for Glyco-core Research (iGCORE), Nagoya University, Nagoya, Japan; Center for One Medicine Innovative Translational Research (COMIT), Nagoya University, Nagoya, Japan; Inamori Research Institute for Science (InaRIS), Kyoto, Japan; Department of Innovative Medicine, Faculty of Medicine, Kindai University, Osaka, Japan

## Abstract

Acute myeloid leukemia (AML) is an aggressive hematologic cancer characterized by proliferation of immature myeloblasts. It shows profound molecular heterogeneity, which has been primarily studied through genetic abnormalities, providing the basis for disease classification, prognostication, and therapeutic choice. However, genetic factors alone may not fully explain AML pathogenesis and diversity, while leaving the role of abnormal epigenome, particularly chromatin state, largely unexplored in a large cohort of patients. Here we show that AML is classified into 16 subgroups with distinct chromatin accessibility profiles based on ATAC-seq in 1,563 AML cases, including novel AML subgroups not previously recognized in conventional genomic classifications. By integrating multi-omics analyses of genome, transcriptome, and major histone marks, we show that these epigenetic subgroups exhibit unique features in clinical presentation, gene mutations, differentiation states, gene expression, and super-enhancer profiles, which are validated across independent cohorts. Single-cell sequencing demonstrates the presence of subgroup-specific ATAC signatures that are shared by all leukemic cells, confirming the definitive role of the epigenome in the ATAC-based classification. Mechanistically, each subgroup is associated with a distinct gene regulatory network centered on key transcription factors, where subgroup-specific super-enhancers play a pivotal role. These ATAC subgroups also have prognostic significance independent of genomic classification, and help reveal unexpected drug sensitivities. In summary, ATAC-based chromatin profiling in this large sample set, combined with multi-omics data, provides new insights into AML pathogenesis beyond genomic profiling and also serves as an invaluable resource for AML research.

## Introduction

Acute myeloid leukemia (AML) is a clinically and molecularly heterogeneous hematologic cancer characterized by dysregulated hematopoietic programs driven by gene mutations, leading to the uncontrolled proliferation of immature myeloblasts in the bone marrow^1^. Over the past decades, advanced genomics studies have substantially improved our understanding of the molecular pathogenesis and heterogeneity of AML by identifying a comprehensive spectrum of major driver mutations and structural variations involved in these dysregulated programs^2–4^. These discoveries have now been incorporated into the latest AML classifications proposed by the World Health Organization (WHO), the International Consensus Classification (ICC) group, and the European LeukemiaNet (ELN), which are widely used for molecular diagnosis, risk stratification, drug discovery, and therapeutic decision-making, including the selection of molecularly targeted therapies^5–8^. However, accumulating evidence suggests that genetic alterations alone may not fully explain AML heterogeneity in terms of gene expression, cellular phenotype, therapeutic response, and prognosis^3,4,9,10^.

The epigenome is a non-genetic mechanism of cellular inheritance and frequently undergoes extensive alterations in cancer through aberrant DNA methylation, chromatin modifications, and higher-order chromatin structure. In fact, the altered epigenome is a hallmark of cancer^11–13^ and serves as another oncogenic mechanism alongside gene mutations, involved in the establishment and maintenance of heterogeneous cancer phenotypes. The cancer epigenome has been most frequently studied through DNA methylation^14–16^, where the methylation analysis predominantly targeted promoter regions using microarray platforms^14^, while non-promoter regions implicated in gene regulation have been less highlighted.

Chromatin is another major component of the epigenome. In concert with DNA methylation and other components, such as transcription factors (TFs) and DNA sequences, chromatin plays a pivotal role in the regulation of gene expression^11^. However, compared to DNA methylation, chromatin has been less intensively studied in cancer using large sample sets because it undergoes many different modifications that frequently need complex experimental platforms for evaluation. Among various chromatin features, we focused on chromatin accessibility. Integrating regulations from different genetic and epigenetic components, chromatin accessibility provides more information than DNA methylation regarding epigenetic regulation of the transcriptional machinery by dictating TF binding and gene expression programs^11,17^. Moreover, chromatin accessibility can be assessed on a genome-wide scale using a simple and scalable method, Assay for Transposase-Accessible Chromatin with high-throughput sequencing (ATAC-seq)^17,18^. Thus, systematic profiling of chromatin accessibility using ATAC-seq is a suitable approach for analyzing the AML epigenome in a large cohort of patients to uncover the epigenetic heterogeneity of AML, although its application has been largely limited to a small number of cases in previous studies^9,18–20^.

In this study, we performed ATAC-seq in a large cohort of AML patients. By integrating multi-layered sequencing data, including known driver mutations, copy number alterations (CNAs), RNA sequencing (RNA-seq), whole-genome sequencing (WGS), chromatin immunoprecipitation sequencing (ChIP-seq), drug sensitivity assays, and single-cell RNA and ATAC sequencing (scRNA/ATAC-seq), we provide an integrated chromatin landscape of AML, elucidate the role of the epigenome in AML pathogenesis and heterogeneity, and explore its impact on clinical presentation and drug sensitivity.

## Chromatin accessibility-based subgroups

In total, we enrolled 1,563 AML patients from independent Swedish and Japanese cohorts and performed ATAC-seq together with targeted-capture sequencing of 331 known/candidate driver genes and >1,000 single nucleotide polymorphism (SNP) sites (for allele-specific copy number detection), RNA-seq, WGS, ChIP-seq, and scRNA/ATAC-seq (Fig. 1a, STables 1-4). Targeted-capture sequencing, complemented with RNA-seq and WGS, reproduced the genomic profile of AML reported in previous studies^2–4^, including all the common driver mutations, gene fusions, and CNAs (ED_Fig. 1a, SFig. 1). Less common mutations involving *MED12*, *PHIP*, *KMT2C*, *MYB*, and *UBTF* were also identified.

**Fig. 1.**
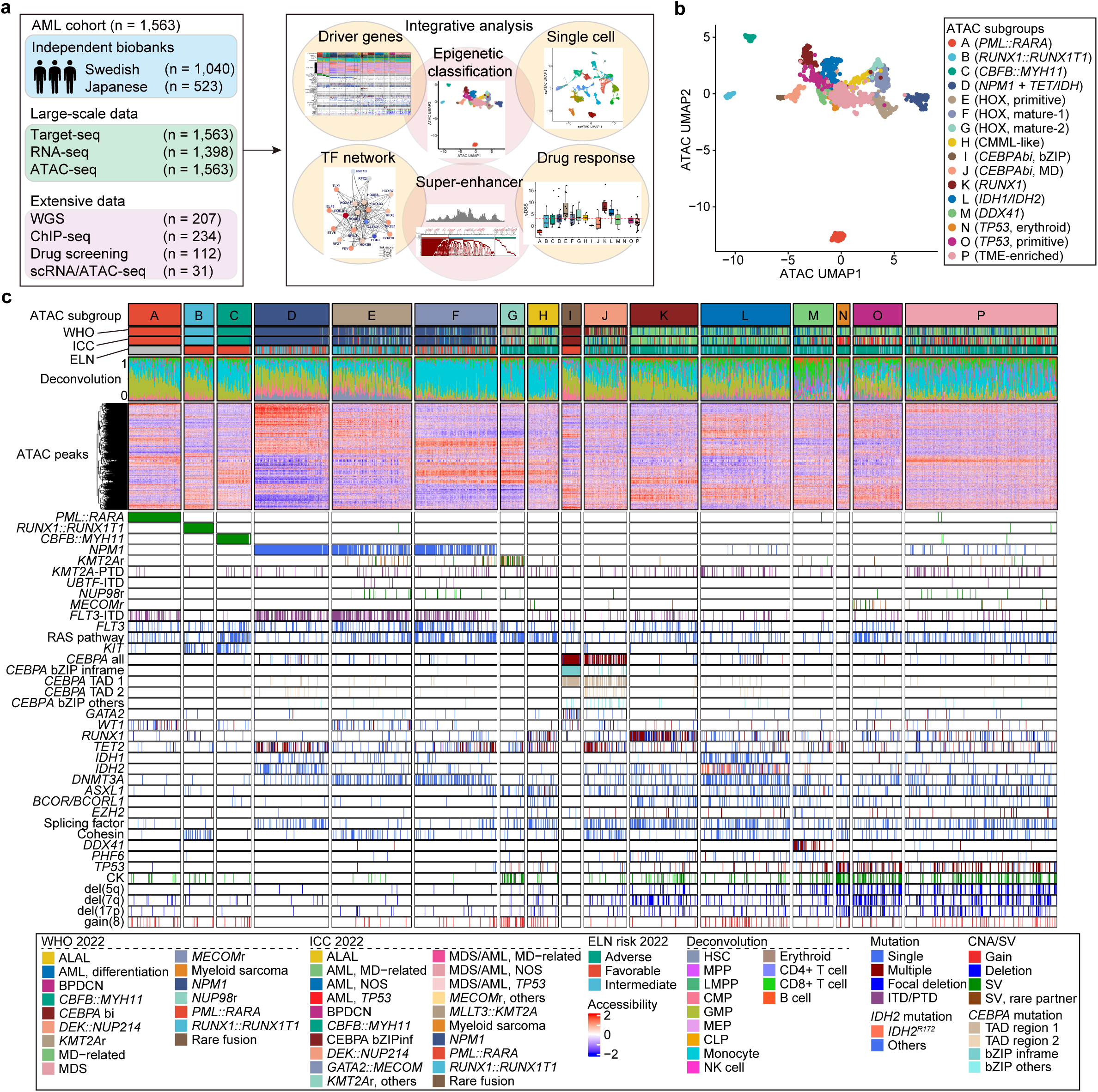
Epigenetic subgroups of AML defined by chromatin accessibility. **a.** Study design and summary of obtained data. **b.** Uniform manifold approximation and projection (UMAP) plot based on ATAC-seq profiles. Each dot represents each patient, and colors indicate ATAC subgroups. MD, myelodysplasia; TME, tumor-microenvironment. **c.** Summary of ATAC subgroups with AML diagnosis, differentiation status inferred by CIBERSORT algorithm using ATAC-seq data (deconvolution), scaled ATAC intensities on 3,000 variable ATAC peaks, and genetic abnormalities. Each column represents each patient. CK, complex karyotype; ALAL, Acute leukemia of ambiguous lineage; BPDCN, blastic plasmacytoid dendritic cell neoplasm; MD, myelodysplasia; MDS, myelodysplastic syndromes; HSC, hematopoietic stem cells; MPP, multipotent progenitor; LMPP, lymphoid-primed multipotent progenitor; CMP, common myeloid progenitor; GMP, granulocyte-monocyte progenitor; MEP, megakaryocyte-erythrocyte progenitor; CLP, common lymphoid progenitor; NK, natural killer; ITD, internal tandem duplication; PTD, partial tandem duplication; CNA, copy number alteration; SV, structural variation; TAD, transactivation domain; bZIP, basic leucine zipper.

In ATAC-seq, we revealed a total of 176,853 recurrent peaks shared between Japanese and Swedish cohorts. These peaks, predominantly located in non-promoter elements, such as introns and intergenic regions, included most of the previously reported ATAC peaks in AML^18^ and also contained many newly identified ones (ED_Fig. 1b). Most ATAC peaks overlapped with one or more ChIP peaks for H3K27ac, SMC1, CTCF, RNA polymerase II, and H3K27me3 that were recurrently detected in ∼200 AML samples (ED_Fig. 1c, SFig. 2), confirming the well-established link between chromatin accessibility and various elements in epigenetic regulation, including promoters, enhancers, and insulators. Although ATAC signals in non-promoter regions were less strong than those in promoters, they showed larger variance between samples (ED_Fig. 1d-e), suggesting that chromatin accessibility in non-promoter elements, such as enhancers, may serve as a clue to understanding epigenetic heterogeneity in AML.

**Fig. 2.**
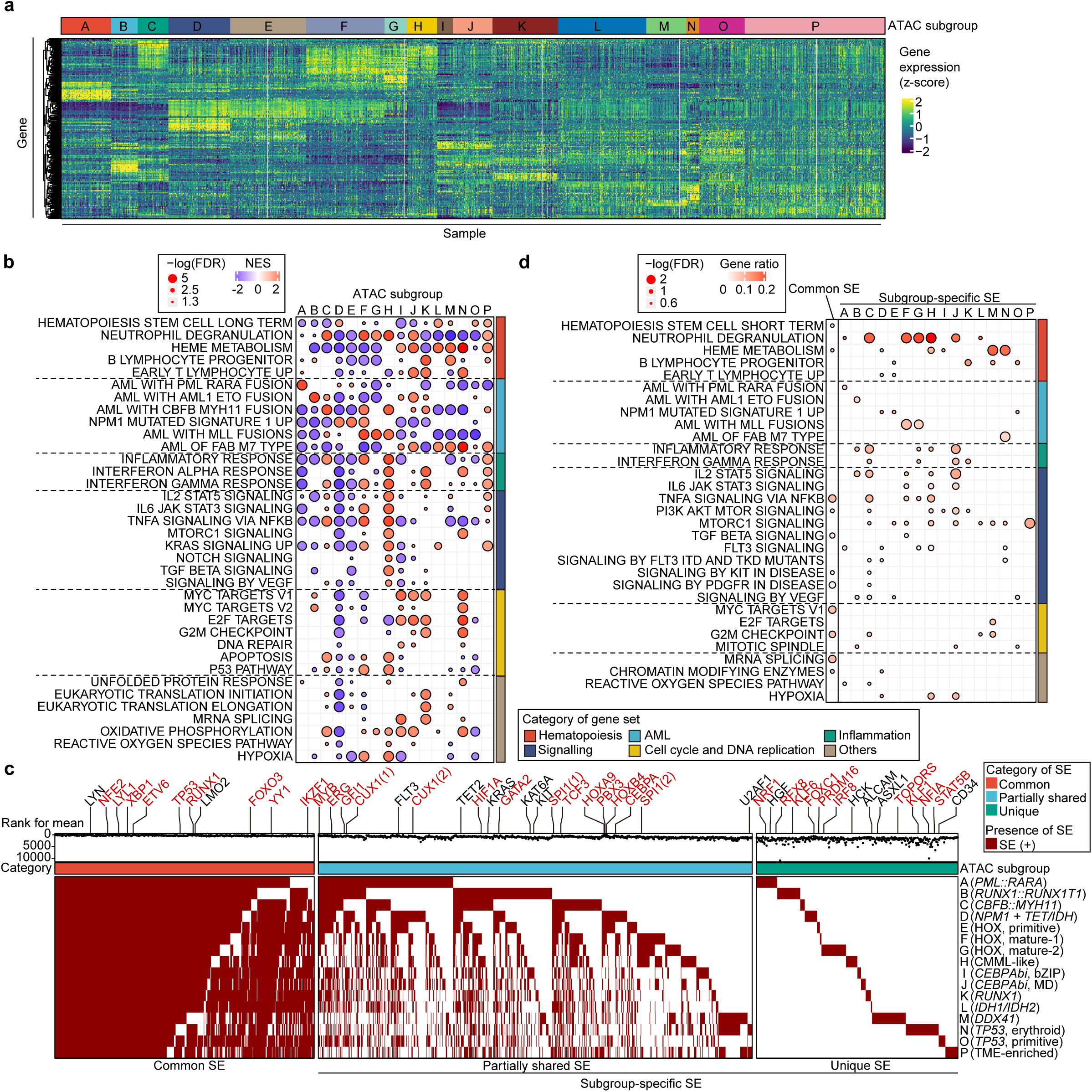
Distinct gene expression and SE profiles across subgroups. **a.** Heatmap for scaled expression of marker genes that were defined by ClaNC algorithm (ED_Fig. 4a) according to ATAC subgroups. Each column represents a sample, and each row represents a gene. **b.** GSEA analysis based on bulk RNA-seq, showing up– and down-regulated gene sets in each subgroup. NES, normalized enrichment score; FDR, false discovery rate. **c.** Distribution of SEs across subgroups, categorized as common, partially shared, and unique SEs. Enhancers are ranked by mean enhancer activities across all AML samples. Representative known oncogenes associated with SEs are indicated, with TFs highlighted in red. **d.** Enriched gene ontologies for identified common and subgroup-specific SEs. Curated gene sets used in **b** and **d** were described in STable 10.

Based on these observations, we sought to reveal epigenetic subgroups of AML using ATAC-based Leiden clustering of 1,563 AML patients and identified 16 subgroups with unique ATAC profiles (Fig. 1b-c). As expected from the close link between the epigenome and differentiation state^18^, each subgroup showed unique lineage commitment, which was revealed by the deconvolution analysis of bulk ATAC-seq data^18,21^ (ED_Fig. 2a-d) and supported by their close correlation with the FAB classification^22^ (ED_Fig. 2e-f). Although primarily characterized by distinct ATAC profiles, these ‘epigenetic’ subgroups showed a unique enrichment of gene mutations and other genetic lesions (Fig. 1c) and significantly overlapped with known genetic subtypes of AML in the WHO and ICC classifications^5,6^ (ED_Fig. 2g-i). However, except for three subgroups defined by common gene fusions, i.e., *PML::RARA* (subgroup A), *RUNX1::RUNX1T1* (subgroup B), and *CBFB::MYH11* (subgroup C), they were not perfectly aligned with existing genetic subgroups, but often showed a complex mapping, wherein a single epigenetic subgroup could be projected onto multiple genetic subgroups in the WHO and ICC classifications, or vice versa (ED_Fig. 2h-i), leading to the identification of novel subtypes not previously defined by the current genomic classification. Notably, the ATAC subgroups overall showed better correlations with leukemia phenotypes than those defined by gene mutations in the WHO and ICC, in several respects, such as gene expression, white blood cell counts, blast percentage, and differentiation profiles (ED_Fig. 2j).

## Characterization of ATAC subgroups

Unique features of ATAC-based classification are highlighted in many newly identified subgroups. For example, most cases in four subgroups (D-G) were classified as *NPM1*-mutated or *KMT2A*-rearranged (*KMT2Ar*) AML in the conventional genomic classifications. This is in line with recent studies suggesting that *NPM1*-mutated AML could be separated into multiple subgroups on the basis of transcriptome and DNA methylation analyses^10,15,16^. However, in the ATAC-based analysis, the separation occurred in a more integrated manner, combining other HOX-related alterations, including *NUP98::x, KAT6A::x,* and *DEK::NUP214* rearrangement, *KMT2A* partial tandem duplication (PTD), and *UBTF* internal tandem duplication (ITD)^23–25^, to generate these four subgroups (D-G) characterized by HOX gene overexpression (ED_Fig 3a-c). Despite the common enrichment of these HOX-related drivers, they showed unique mutation patterns and cellular differentiation. For example, *NPM1* and *FLT3*-ITD mutations were highly condensed in subgroups D-F, but almost absent in subgroup G, which was instead enriched for *KMT2Ar* together with complex karyotype (CK) and gain of chromosome 8. Cases in subgroup D were discriminated from other *NPM1*-enriched subgroups by the presence of mutually exclusive *TET2*, *IDH1*, and *IDH2* mutations in virtually all cases, a high frequency of *SRSF2* mutations, and a paucity of *DNMT3A* mutations, particularly the R882 hotspot mutations (ED_Fig. 3d-f, SFig. 3). Subgroup E frequently had *WT1* and cohesin (*STAG2*, *RAD21*, *SMC1A*, and *SMC3*) mutations. Subgroups D and E exhibited immature phenotypes with an increased hematopoietic stem cells and progenitor (HSPC) component, whereas subgroups F and G, largely classified as FAB M4 and M5 subtypes, had markedly increased monocytic components, irrespective of the type of associated HOX-related drivers (ED_Fig. 2a-b,e). Marked monocytic differentiation was also seen in subgroup H. Notably, this subgroup had a characteristic co-mutation pattern that is reminiscent of chronic myelomonocytic leukemia (CMML), with frequent mutations in *TET2*, *RUNX1, ASXL1, SRSF2*, as well as *RAS* pathway mutations, suggesting that they might have been derived from or represent a continuum of CMML^26,27^(ED_Fig. 3a, SFig. 3).

**Fig. 3.**
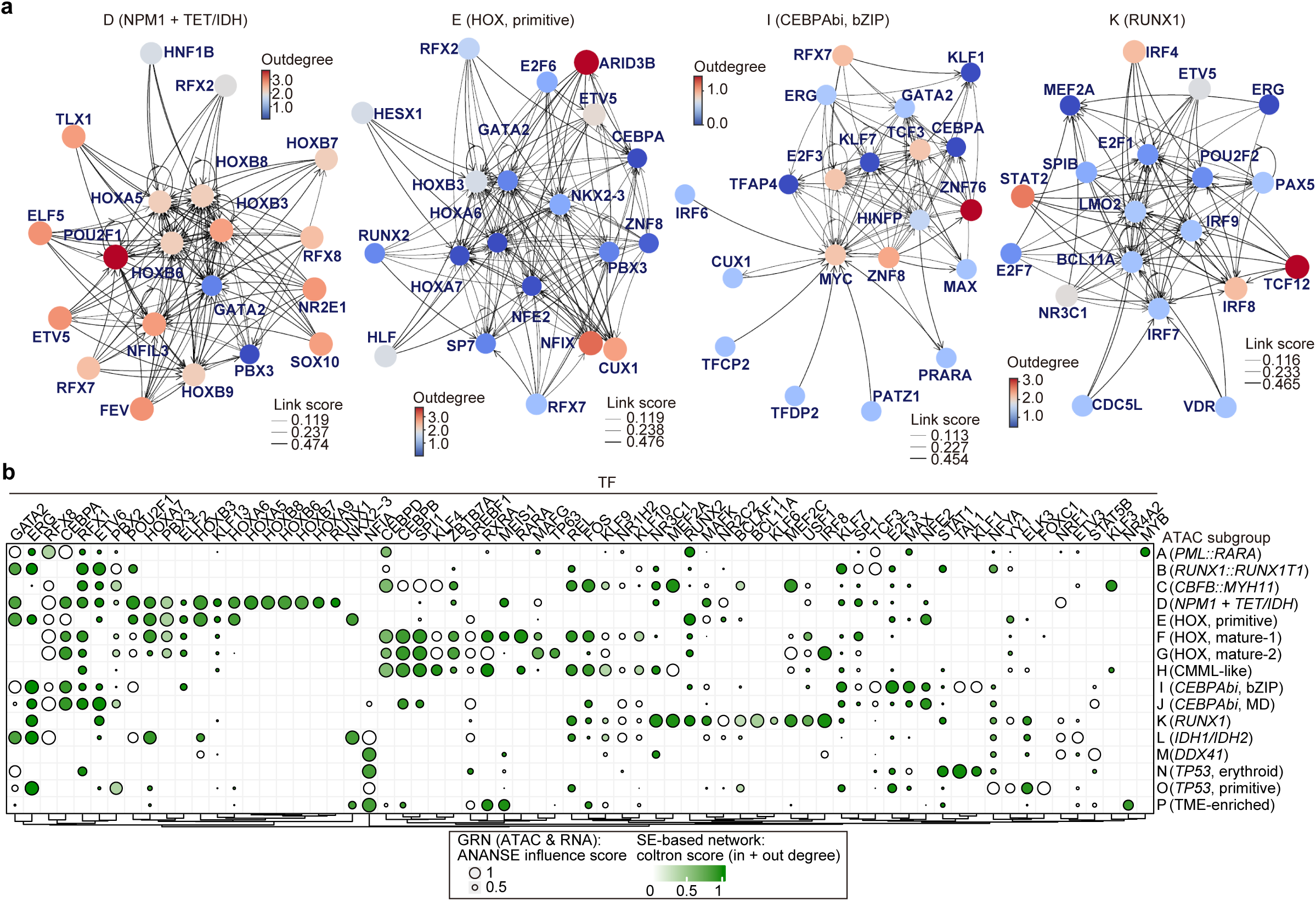
Subgroup-specific GRNs driven by SE-regulated TFs. **a.** GRNs centered on TFs specific to indicated subgroups, generated using ANANSE software_38_ (ED_Fig. 6a). Top 20 TFs that most significantly contributed to subgroup-specific gene expression (influence scores) in each ATAC subgroup are shown. Colors indicate the number of TF genes each TF regulated (outdegrees). Line width represents the strength of expressional regulation (linkscores) between two TFs. **b.** Heatmap summarizing activities of TFs that were identified in both ANANSE and coltron analysis. Size indicates influence scores for each TF, calculated by ANANSE software_38_ (ED_Fig. 6a). Color indicates the importance of each TF in SE-based TF networks (coltron score), generated by the coltron software_39_, determined by the sum of in-degree and out-degree in the networks (ED_Fig. 6c).

Subgroups I and J were characterized by an enrichment of biallelic *CEBPA* mutations. Of these, largely conformed to the ICC subtype defined by in-frame basic leucine zipper (bZIP) *CEBPA* mutations^5^, subgroup I was characterized by the combination of bZIP and transactivation domain (TAD) region I mutations in most cases, along with common mutations in *GATA2* and *WT1* (ED_Fig. 3a,g-h). By contrast, the enrichment of biallelic *CEBPA* mutations in subgroup J was less conspicuous, accounting for only 59% of the cases, where the biallelic *CEBPA* mutations appeared in other combinations than the typical bZIP and TAD region I mutations (ED_Fig. 3g-h). However, otherwise this subgroup was uniformly accompanied by mutations in *TET2* and MDS-related mutations such as those in *ASXL1*, cohesin, and splicing factor genes (ED_Fig. 3a, SFig. 3).

Subgroup K exhibited a strong enrichment of *RUNX1* mutations together with MDS-related alterations, particularly splicing factor mutations as well as del(7q) and CK. Subgroup L was characterized by mutually exclusive *IDH1* and *IDH2* mutations with frequent *IDH2^R172^* variants (ED_Fig. 3i). Subgroup M exhibited an enrichment of *DDX41* mutations and erythroid differentiation (ED_Fig. 2a,3a). The remaining subgroups (N-P) were enriched for *TP53* mutations (ED_Fig. 3a), of which subgroups N and O were highly skewed to erythroid and megakaryocytic-erythroid progenitor (MEP) lineages and HSPCs, respectively (ED_Fig. 2a,d-e), whereas Subgroup P showed balanced contributions from various hematopoietic lineages and lower blast percentages in the bone marrow.

## Gene expression and super-enhancers

We next investigated other molecular features of epigenetic subgroups by integrating RNA-seq, ATAC-seq, and ChIP-seq. Each epigenetic subgroup exhibited unique patterns of gene expression (Fig. 2a), which were enriched in specific biological pathways (Fig. 2b), suggesting distinct gene expression programs. For example, subgroups D-G were characterized by HOXA gene overexpression, yet each showed subgroup-specific gene expression and pathway enrichment patterns, such as the downregulation of cell cycle/DNA replication (D) and TGFβ signaling (E), as well as the upregulation of inflammatory response (F) and neutrophil degranulation (F and G) (Fig. 2b, SFig. 4). Subgroups C, F, and H, all showing prominent monocytic differentiation, shared upregulated TNFα signaling, IFNγ response, and inflammatory response, which were absent in subgroup G. Subgroups I and J had upregulated expression of genes related to MYC and E2F targets, hem metabolism, and oxidative phosphorylation, while other pathways, such as inflammatory responses and hypoxia, were selectively downregulated in subgroup I.

**Fig. 4.**
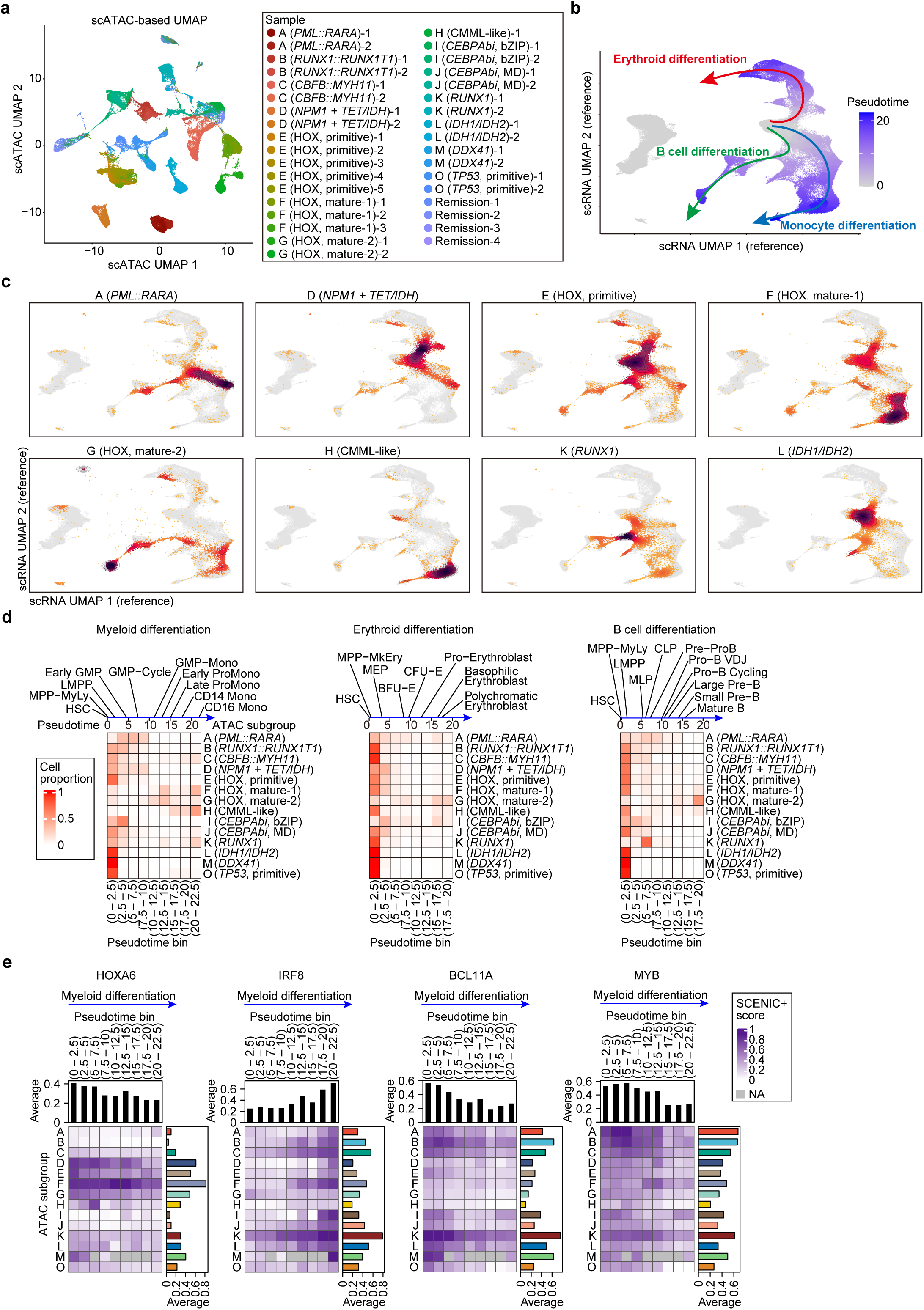
Epigenetic signatures, differentiation states, and TF dynamics at the single-cell level. **a.** UMAP plot based on scATAC-seq profiles for all sequenced samples. Each dot represents one cell, and colors indicate different patient samples. Each sample is labeled with its group and sample number. **b.** Scheme for reference-based pseudotime analysis. Color indicates pseudotime. **c.** Density plot showing the distribution of AML cells mapped from each ATAC subgroup onto the reference scRNA-seq UMAP. **d.** Proportion of cells distributed across pseudotime bins along differentiation trajectories. The mean pseudotime for representative cell types in the normal hematopoiesis (ED_Fig. 7d) is indicated at the top. **e.** TF activity as inferred by SCENIC+ score (gene-based Area Under the Curve (AUC) in SCENIC+ software_47_ (ED_Fig. 8a)) in each pseudotime bin along the myeloid differentiation trajectory. The top barplot shows the average SCENIC+ score per bin, and the right barplot displays the average SCENIC+ score per subgroup.

Given their distinct gene expression profiles across subgroups, we built an expression-based prediction model for ATAC subgroups using the Classification to Nearest Centroids (ClaNC) algorithm^28^ (METHODS, ED_Fig. 4a-b). Using this model, we predicted ATAC subgroups in a total of 1,079 samples from four external adult AML cohorts^2,10,29,30^, successfully recapitulating the enrichment of gene mutations and clinical features characteristic of each subgroup, thereby confirming the robustness and validity of our ATAC-based classification (ED_Fig. 4c). In addition, the prediction model also enabled us to analyze DNA methylation across ATAC subgroups, using the DNA methylation data available in the BeatAML cohort^15^. As expected, each ATAC subgroup had a unique methylation profile (ED_Fig. 4d). Notably, the ATAC signals and DNA methylation levels at corresponding CpG sites exhibited almost perfect mirror images across subgroups, highlighting a strong link between chromatin accessibility and DNA methylation (ED_Fig. 4e).

To further understand the molecular basis of the unique gene expression in ATAC subgroups, we next analyzed subgroup-specific super-enhancers (SEs), as SEs are known to define cellular identity and drive oncogene expression^31,32^. For this purpose, we first identified all enhancers recurrently found in a total of 234 AML samples using H3K27ac ChIP-seq data, ranked them by average H3K27ac signals in each subgroup, and identified the top 750 enhancers as SEs in that subgroup. After combining the detected SEs from all subgroups, SEs were classified as common (n=498, ≥12 subgroups), partially shared (n=833, detected in 2–11 subgroups), or unique (n=387, found in a single subgroup) SEs (Fig. 2c, STable 6). Common SEs showed a greater overlap with SEs detected across various tissues and cell types^32^, particularly in hematopoietic cells, such as CD34-positive HSPCs, whereas subgroup-specific SEs were more AML-specific (SFig. 5). Common and subgroup-specific SEs were linked to different sets of genes. Common SEs were implicated in the regulation of oncogenic TFs, such as *ETV6*, *TP53*, *RUNX1*, and *LMO2*, partially shared SEs were mapped to genes such as *MYB*, *ERG*, *FLT3*, *HOXA9*, *CEBPA*, and *SPI1* (ED_Fig. 5a), and unique SEs targeted known oncogenes, such as *RFX8* and *HGF* in subgroup A, *FOXC1* in subgroup D, *PRDM16* in subgroup E, *IRF8* in subgroup G, *KLF1* and *NFIA* in subgroup N, and *CD34* in subgroup O (ED_Fig. 5b-d). These subgroup-specific SEs were shown to tightly correlate with accessible chromatin and Pol II binding with variable cohesin signals (ED_Fig. 5a-c). Gene ontology analysis revealed distinct associations of genes regulated by common and subgroup-specific SEs (Fig. 2d). Common SEs were associated with genes enriched for TNFα signaling, MYC targets, G2M checkpoint, and mRNA splicing, while subgroup-specific SEs often showed an enrichment for the gene sets reported for the related AML subtypes; AML with *PML::RARA*, *RUNX1::RUNX1T1*, *NPM1* mutation, and *KMT2Ar* corresponding to subgroups A, B, D-E, and F-G, respectively. Other gene sets enriched for subgroup-specific SEs included those implicated in neutrophil degranulation and TNFα signaling (C, F, G, and H), aligning with overexpression of these genes in corresponding subgroups (Fig. 2b). To understand the role of these SEs in subgroup-specific AML pathogenesis, we evaluated their association with lineage-specific gene expression (METHODS and ED_Fig. 5e). As expected, these subgroup-specific SEs were less associated with lymphoid differentiation. SEs in M-N and F-H subgroups were associated with high expression of genes in erythroid and myelomonocytic cells, respectively, which is consistent with their lineage contributions from the corresponding cell lineages (ED_Fig. 5e,2a). In subgroups D and O, SEs were associated with gene expression related to hematopoietic stem cells (HSCs), suggesting the role of the HSC-related program in these subgroups. Taken together, these results suggest that each ATAC subgroup was characterized by the unique SE profile that was tightly correlated with gene expression and cell differentiation programs.

**Fig. 5.**
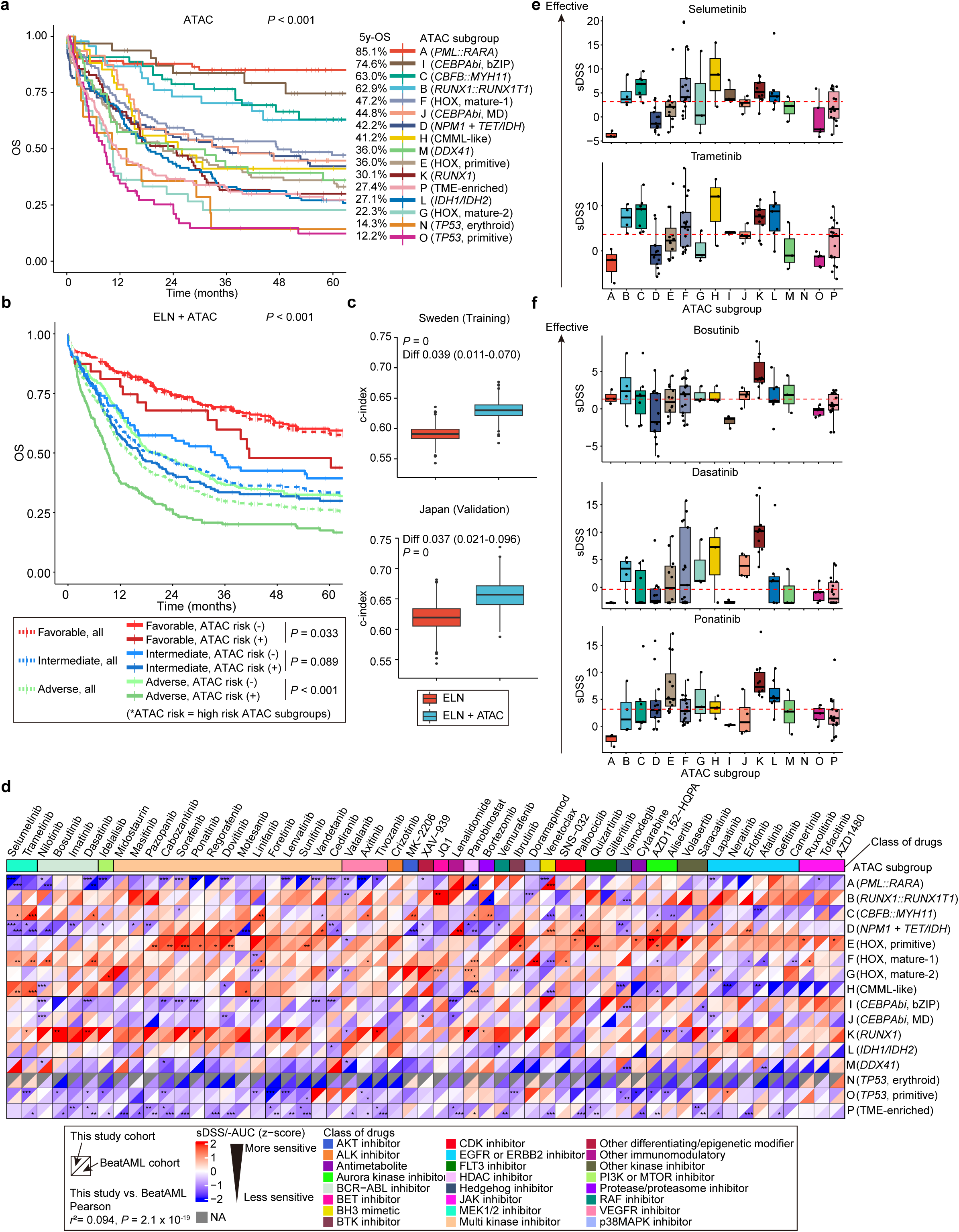
Distinct prognoses and drug sensitivities across subgroups. a,b. Kaplan–Meier survival curves for OS of patients who received intensive chemotherapy, categorized by ATAC subgroups (**a**) and combined model of ELN and high risk ATAC subgroups (**b**). *P* values were calculated using the log-rank test. High-risk ATAC subgroups for each ELN risk group in panel **b** were determined as shown in ED_Fig. 9b: ATAC subgroup E for ELN favorable, subgroups E, F, G, K, L, and M for ELN intermediate, and subgroups D, E, G, O, and P for ELN adverse risk. **c.** Estimates of the concordance index (c-index) derived from ELN and ELN combined with ATAC risk subgroups, using bootstrapping 1,000 times. The combined model was trained in the training (Swedish) cohort and evaluated in both training and validation (Japan) cohorts. *P*-value was computed using 1,000 bootstrap replicates, defined as the fraction of replicates in which the c-index difference (ELN+ATAC minus ELN) was ≤ 0. **d.** Heatmap summarizing sensitivities to each drug across subgroups. The upper left and bottom right triangles in each box indicate normalized selective drug sensitivity score (sDSS) for our dataset and normalized area under the curve (AUC) for BeatAML dataset, respectively, where red indicates higher drug sensitivities. *P* values were calculated by Student’s t-test, corrected for multiple testing by Benjamini-Hochberg method. *FDR <0.05; ** FDR <0.01; *** FDR <0.001. Pearson’s correlation between our and BeatAML dataset was calculated for all boxes in the heatmap. NA, not available. **e,f.** Sensitivities to MEK (**e**) and ABL inhibitors (**f**) for each ATAC subgroup. y-axis shows sDSS, in which higher values indicate increased sensitivity and positive values indicate effectiveness in AML cells compared to control CD34 positive cells. Dashed red line indicates median sDSS value across all patients.

## Gene regulatory mechanism

Given that ATAC subgroups had distinct profiles of gene expression and SEs, we wanted to understand the underlying molecular mechanisms of these findings. For this purpose, we reasoned that deregulated TFs play key roles in the pathogenesis of AML through altered gene expression programs^33^ and analyzed subgroup-specific gene regulatory networks (GRNs) centered on TFs using the ANANSE algorithm^34^. By integrating ATAC-seq and RNA-seq, this algorithm infers the TFs most significantly contributing to subgroup-specific gene expression for each ATAC subgroup by calculating “TF influence scores”. It also calculates the number of genes they regulate (outdegrees) and the strength of the regulation (linkscores) (METHODS and ED_Fig. 6a). Overall, each ATAC subgroup was associated with a unique GRN of TFs responsible for the subgroup-specific gene expression (Fig. 3a, ED_Fig. 6b).

Since TFs defining cell identity frequently occupy SEs and form regulatory networks to determine gene expression programs^31,32^, we next sought to correlate key TFs identified in ANANSE analyses with subgroup-specific SEs. To this end, we built another network of TFs that were regulated by SEs using the coltron algorithm^35^. In the coltron-based network, TF–TF connections were inferred from motif occurrences within SEs that regulate each TF, capturing both out-degree (regulatory targets) and in-degree (upstream regulators) (see Methods and ED_Fig. 6c). Notably, the influence score of the key TFs in the ANANSE-based network (expressed as the size of circles) significantly correlated with the sum of in– and out-degrees in the coltron analysis (expressed as color gradient) (Fig. 3b, ED_Fig. 6d), suggesting the role of subgroup-specific SEs in the distinct gene regulatory programs across ATAC subgroups via key TFs. For example, HOXA family TFs and E2F3, key TFs in GRNs in subgroups D-E and subgroup I, respectively, were typical TFs regulated by SEs in respective subgroups (Fig. 3b). BCL11A and IRF family TFs (subgroup K) and SPI1 and CEBPA family TFs (subgroups C, F, G, and H) were other examples of SE-associated TFs implicated in B-cell development^36,37^ and monocytic differentiation^38,39^, respectively, suggesting that the regulation of these TFs via subgroup-specific SEs might explain the increased B-cell and monocytic components found in the corresponding subgroups (ED_Fig. 2a). Taken together, these findings support the hypothesis that each ATAC subgroup is regulated by a distinct set of key TFs, which are often regulated by SEs specific to each subgroup.

## Single-cell profiling of ATAC subgroups

We next performed multi-omics scRNA/ATAC-seq analysis to validate the findings from bulk sample analysis in greater detail and obtain further insights into the role of TFs in the regulation of gene expression in ATAC subgroups. We analyzed 233,713 mononuclear cells from 31 AML samples across 14 ATAC subgroups along with four complete remission samples as normal controls (Fig. 4a, ED_Fig. 7a, SFig. 6-7, STable 7). In scATAC-seq-based analysis, the leukemic cells from each AML sample tended to be clustered into a single, well-defined group, which was distinct from the residual normal cells co-clustered with remission-derived cells (Fig. 4a, ED_Fig. 7b-c). Notably, samples belonging to the same ATAC subgroup were mostly co-clustered, despite being derived from different patients (ED_Fig. 7c). In particular, five subgroup E samples harboring distinct HOX-related driver mutations—including *NPM1* mutations and (n=3) *NUP98* rearrangements (n=2)—were predominantly assigned to the same cluster. Although some subgroups (E and F, and I and J) were not completely separated from each other, these observations indicate that all leukemic cells in each subgroup shared a distinct ATAC signature regardless of their differentiation states, which uniquely clustered them together. A similar co-clustering pattern was also observed in scRNA-seq analysis, confirming again the strong impact of the epigenome on gene expression (ED_Fig. 7a, SFig. 7).

Despite the unique clustering across samples, deconvolution analysis based on bulk ATAC-seq data suggests that each leukemic sample may consist of heterogeneous populations with various cell lineages (ED_Fig. 2a). Thus, we investigated the differentiation profile of leukemic cells at the single cell level. By projecting scRNA-seq data of each ATAC subgroup to a reference trajectory of normal bone marrow cell differentiation generated by scRNA-seq-based imputation^40^, we delineated unique lineage commitment and differentiation dynamics in ATAC subgroups that mimicked normal blood cell differentiation (Fig. 4b-d, ED_Fig. 7d-g). The differentiation profiles were largely concordant between samples within the same ATAC subgroup, also supporting the biological significance of the ATAC-based clustering (SFig. 8). For example, seven samples from HOX-related subgroups (D-F) shared common *NPM1* mutations yet showed distinct differentiation profiles. Three samples from subgroup E showed a maturation block at the HSC stage, two from subgroup D were arrested at the GMP stage, and the remaining two from subgroup F comprised both progenitor and mature cell populations. Of note, two *NUP98*-mutated samples from subgroup E also displayed differentiation profiles similar to those found in the *NPM1*-mutated samples from the same subgroup. Although subgroups F-H showed prominent monocytic components in bulk sample analysis (ED_Fig. 2a), single-cell analysis revealed that mature monocyte-like cells were predominant in subgroup H, whereas samples from subgroups F and G contained more immature, promonocyte-like cells (Fig. 4c-d). Of interest, subgroup K samples were blocked at the common lymphoid progenitor stage along the B-cell trajectory and showed a differentiation block toward the myeloid and erythroid lineages, indicating that a commitment to lymphoid lineages is a unique feature of the pathogenesis of this subtype (Fig. 4c-d).

We then sought to evaluate the distinct TF activities across ATAC subgroups leveraging multiome scRNA/ATAC-seq data (ED_Fig. 8a). Using the SCENIC+ algorithm^41^, we inferred key TFs enriched in each subgroup, which were significantly overlapped with those found in bulk-based analysis (ANANSE) and also in coltron analysis (ED_Fig. 8b-d). We then investigated the activity of key TFs along the myeloid differentiation trajectory by combining SCENIC+ and pseudotime analysis, focusing on those reproducibly found in the three analyses (ANANSE, coltron, and SCENIC+) (Fig. 4e, ED_Fig. 8e). For example, HOXA6 was consistently activated throughout the myeloid differentiation trajectory in HOX-related subgroups (D-G) but peaked at different stages according to subgroups (Fig. 4e). By contrast, although consistently activated along the myeloid trajectory in subgroup K, IRF8 and BCL11A were differentially peaked in mature and immature stages, respectively. Another example of interest was MYB, which showed a characteristic activation peak synchronized with a differentiation arrest at the GMP stage in subgroup A. These findings suggest that the activation of key TFs at critical stages along the differentiation trajectory may explain the subgroup-specific leukemogenic mechanisms.

## Clinical features and drug sensitivity

Finally, we investigated whether the ATAC subgroup correlated with patient clinical characteristics and predicted drug sensitivity, using the BeatAML cohort for validation. Each ATAC subgroup exhibited unique clinical features, including age, white blood cell count, blast percentage, and the prior history of myeloid disease and chemotherapy, which were largely similar in the BeatAML cohort (SFig. 9). We next assessed the prognostic impact of ATAC subgroups by evaluating overall survival (OS) in patients who received intensive chemotherapy. OS differed significantly across ATAC subgroups in both AML cohorts, with similar prognostic impacts observed (Fig. 5a, SFig. 9d). Specifically, subgroups A, I, C, and B were associated with favorable prognosis, whereas subgroups L, G, N, and O showed poor outcomes. To determine whether the ATAC-based classification provides prognostic value independent of the established ELN risk stratification, we evaluated the effect of each ATAC subgroup on OS within each ELN risk category using multivariable analyses that included clinical factors. As shown in Fig. 5b, OS within each ELN risk category was significantly influenced by one or more ATAC subgroups (ED_Fig. 9a-b). The effect of these ATAC subgroups was demonstrated independently in the Japanese and Swedish cohorts, where the inclusion of ATAC risk subgroups increased the concordance index (Fig. 5c).

We next investigated the impact of ATAC subgroups on drug sensitivity obtained from 112 samples tested against 250 drugs^42,43^. We also analyzed the drug sensitivity data from the BeatAML study (*n* = 569, for 166 drugs)^29^ based on the ATAC subgroups inferred in ED_Fig. 4. The results were combined and summarized for the 55 drugs tested in both cohorts (Fig. 5d). Reproducible drug sensitivities in both cohorts were identified for multiple drugs and ATAC subgroups. Some of these drug-subgroup correlations were strictly determined by the target mutations. For example, enriched for *FLT3* mutations, subgroup E exhibited high sensitivity to FLT3 inhibitors, including quizartinib and giltertinib, where the efficacy was observed only for the *FLT3*-mutated, but not for unmutated samples (Fig. 5d, ED_Fig. 10a-b). By contrast, the sensitivity to MEK1/2 inhibitors, such as selumetinib and trametinib, was observed in subgroups C, F, and H, not only in samples carrying RAS pathway mutations enriched in these subgroups but also in those lacking known RAS pathway mutations (Fig. 5d-e, ED_Fig. 10c). This suggests that drug sensitivities may be determined not solely by gene mutations but also by epigenetic mechanisms, which could be captured by ATAC subgroups.

The ATAC subgroups also revealed drug sensitivities that were not previously predicted by gene mutation profiles, which was exemplified by an unexpected sensitivity of subgroup K to ABL inhibitors (Fig. 5f, ED_Fig. 10d). Because these samples were sensitive to multiple ABL inhibitors (e.g., imatinib, nilotinib, dasatinib, bosutinib, and ponatinib), this sensitivity was most likely explained by the inhibition of ABL kinases, suggesting the key role of activated ABL kinases in the pathogenesis of this ATAC subgroup. As already seen in ED_Fig. 3a, subgroup K was characterized by a strong enrichment of *RUNX1* mutations. However, *RUNX1* mutations alone could not fully explain the sensitivity to ABL inhibitors in this subgroup, since *RUNX1*-mutated samples in other ATAC subgroups did not exhibit similar sensitivities (ED_Fig. 10e). Thus, taken together, these findings highlight the role of ATAC subgroups in the discovery and prediction of drug sensitivity.

## Discussion

We conducted an integrated analysis of the AML epigenome, generating a large dataset comprising ATAC-seq, ChIP-seq, and multi-omics scRNA/ATAC-seq data (Fig. 1a). To our knowledge, this represents the most comprehensive epigenomic dataset for a single cancer type. By integrating gene mutation, transcriptome, and drug sensitivity data, it serves as an invaluable resource for investigating AML pathogenesis and therapeutic strategies.

A central finding of our study is the identification of distinct epigenetic subgroups in AML using ATAC-based clustering, where each subgroup is associated with unique clinical and molecular features. Given that most of these subgroups are newly identified (e.g., H, K, and L) or represent a reorganization of multiple subtypes (e.g., D-G, I-J, M-P) in conventional genomic classifications, the ATAC-based classification reveals epigenetic heterogeneity of AML, offering a novel framework for understanding AML pathogenesis from a perspective distinct from the conventional genomic classification. Meanwhile, the compromised performance of clustering is suspected for some subgroups (e.g., O and P), suggesting that a larger cohort size is required to improve the resolution of subgroup classification.

scATAC-seq demonstrates that the identification of these heterogeneous epigenetic subgroups in bulk sample analysis is enabled by the presence of subgroup-specific, heritable chromatin accessibility profiles shared by all leukemic cells. However, the exact mechanisms by which these subgroup-specific epigenetic profiles are established in the most recent common leukemic ancestor (MRCA) are not fully understood. While the impact of many driver mutations on the epigenome—through DNA methylation and chromatin modifications, either directly or indirectly—is well anticipated, the role of primary alterations of the epigenome remains unclear. An important issue that remains to be elucidated is how such primary epigenetic alterations or ‘epi-mutations’ can directly contribute to the positive selection of the MRCA to establish the highly heterogeneous leukemic population.

The newly identified ATAC subgroups can be effectively used to investigate AML pathogenesis by combining multi-layered epigenetic and transcriptional analyses. In this study, these analyses were used to uncover subgroup-specific gene expression profiles and to comprehensively identify SEs in AML. These findings were then integrated with ATAC-seq data to delineate subgroup-specific gene regulatory mechanisms. Our findings contribute to understanding the heterogeneous mechanisms of leukemogenesis, highlighting the role of TFs and SEs in dysregulated gene expression across ATAC subgroups, although further functional validation is required. The strong epigenome-transcriptome correlations enabled gene expression-based inference of ATAC subgroups, validating the findings from bulk sequencing in independent external cohorts. This also provides the foundation for constructing a classifier to infer ATAC subgroups, facilitating their application in clinical settings.

Our ATAC-based epigenetic classification also has important clinical implications. ATAC subgroups are significantly associated with overall survival and provide prognostic value independent of conventional models such as the ELN risk classification^7^. Integrating ATAC subgroups into prognostic models could enhance AML risk stratification. Last but not least, ATAC subgroups are associated with distinct drug sensitivity profiles, and, as such, enable the identification of novel drug susceptibilities that are not predicted by gene mutations alone. Moreover, they help refine personalized therapeutic strategies by guiding drug combinations tailored to individual patients.

## Methods

### Patient samples

This study was reviewed and approved by the institutional ethics committees of the participating institutions or biobanks, and was performed in accordance with the Declaration of Helsinki. Informed consent was obtained from all participants at participating instituions. Consecutively diagnosed patients were included and no statistical methods were used to determine the sample size. A total of 1,563 patients were enrolled from the Swedish and Japanese cohorts, who were diagnosed with AML and related neoplasms^7^, which sometimes exhibit a continuous and unclear boundary with AML, including 16 with acute leukemia with ambiguous lineage (ALAL), 11 with myelodysplastic syndromes (MDS), 6 with myeloid sarcoma (MS), and 1 with blastic plasmacytoid dendritic cell neoplasm (BPDCN) (STables 1-2). Detailed clinical annotations, including patient information, diagnosis, clinical laboratory values, treatments, and outcomes were collected from electronic medical records and from the Swedish AML Registry. Tumor samples, such as bone marrow or peripheral blood, and matched control buccal samples, were obtained from patients. We also analyzed normal bone marrow samples from 25 individuals without hematological malignancies who underwent hip joint replacement surgery, serving as controls for the ATAC-seq analysis. Bone marrow and peripheral blood cells were isolated, subsequently subjected to erythrolysis or mononuclear cell (MNC) isolation by Ficoll gradient centrifugation, resuspended in CELLBANKER 1 solution (Nippon Zenyaku Kogyo), and cryopreserved in liquid nitrogen. Genomic DNA was extracted using the QIAamp DNA mini kit (Qiagen), Gentra PureGene kit (Qiagen), or Maxwell® RSC Genomic DNA Kit (Promega). RNA was extracted using RNeasy Mini Kit (Qiagen) or Maxwell® RSC simplyRNA Tissue Kit (Promega). A summary of this cohort, including diagnosis, ATAC subgroup, AML classifications, driver genes, and available multi-omics data, is provided in STable 3.

### Targeted-capture sequencing

Targeted-capture sequencing was performed using the SureSelect custom kit (Agilent Technologies) with an in-house gene panel including 331 known AML driver genes (STable 4) and an additional 1,158–1,317 probes for copy number detection. Captured targets were sequenced using the NovaSeq 6000 (Illumina) or DNBSEQ G400 (MGI) instrument with a 150-bp paired-end read protocol.

Sequence alignment and mutation calling were performed using the hg19 reference genome and Genomon pipeline (https://github.com/Genomon-Project)^45–48^. The called variants were further filtered by assessing the oncogenicity of variants based on an in-house curation program^45–48^ which utilizes the COSMIC database (v96), an in-house blacklist of error calls, and public SNPs databases, such as 1000 Genomes Project (October 2014 release), NCBI dbSNP build 138, National Heart, Lung, and Blood Institute (NHLBI) Exome Sequencing Project (ESP) 6500, Human Genetic Variation Database (HGVD), and our in-house dataset.

### Copy number analysis

SNP probes included in the target bait for targeted-capture sequencing were utilized to allow for detection of copy number changes and allelic imbalances^45–48^. This CNACS program is available at https://github.com/papaemmelab/toil_cnacs. Total copy number (TCN) of 2.22 or larger was defined as gains, and TCN less than 1.88 was assumed as losses. Copy number neutral loss of heterozygosity was called with B-allele frequency <0.90 and TCN between 1.88 and 2.22. Arm level changes were called for regions with the total length >1 Mbp. Detected copy number changes were manually curated.

### Structural variation analysis

Structural variations (SVs) were detected using the Genomon SV pipeline^49^, which utilizes both breakpoint-containing junction read pairs and improperly aligned read pairs. Detected putative SVs were filtered by removing (i) those with <4 supporting tumor reads and <10 supporting tumor/normal reads and (ii) those present in control normal samples, whose breakpoints were manually inspected using the Integrative Genomics Viewer (IGV).

### WGS

WGS data was obtained as part of a large-scale WGS project for cancers in Japan (Genome research in CAncers and Rare Diseases (G-CARD)). Sequencing library was generated using the Illumina DNA PCR-Free Prep Tagmentation kit according to the manufacturer’s protocol, and sequenced using NovaSeq 6000 (Illumina) instrument with a 150-bp paired-end read protocol at a targeted depth of 100x for tumors and 30x for normal controls. The reads were aligned to the human hg38 reference genome by parabricks. SNV calling was performed using the Mutect2 pipeline (GATK version 4.5.0.0) in tumor with matched normal mode, with a variant allele frequency threshold of “normal ≤ 0.3” and “TLOD ≥ 20”. Variants listed in germline databases, such as gnomAD (v3.1.2), 1000 Genomes Project, and Tohoku Medical Megabank Organization (ToMMo), with a minor allele frequency of 0.001 or higher, were removed. Structural variants were called using GRIDSS v2.12.0 and Genomon SV v0.8.0. Filtered outputs from GRIDSS (’FILTER = PASS, QUAL ≥ 500, AS > 0, or RAS > 0’) and Genomon SV (’overhang ≥ 150’) were combined to create their union.

### Bulk RNA-seq experiments and analysis

Libraries for RNA-seq were prepared using the NEBNext Single Cell/Low Input RNA Library Prep Kit for Illumina (New England BioLabs) and were subjected to sequencing using NovaSeq 6000 instrument (Illumina) with a paired-end protocol. The sequencing reads were preprocessed by fastp^50^ with ‘--detect_adapter_for_pe –q 15 –n 10 –u 40’ parameters, and then aligned to the reference human hg19 genome using STAR^51^. Reads on each hg19 UCSC gene were counted with featureCounts^52^. The quality of sequencing data was assessed using mapping and count statistics. Samples were excluded from the analysis if any of the following criteria were met: ‘uniquely_mapped_percent’ (STAR) <30%, ‘percent_assigned’ (featureCounts) <30%, or ‘assigned’ (featureCounts) <3M. The edgeR package^53^ was used to normalize read counts and calculate counts per million (CPM) values of genes. Genes expressed at >1 CPM count in two or more samples and located on the autosomal chromosomes were kept and used for downstream analysis. To correct batch effects between different cohorts or dataset, ‘removeBatchEffect’ command from the limma package^54^ was used. Differentially expressed genes (DEGs) for each subgroup were identified using the eBayes test in limma through a one-versus-rest comparison, with thresholds of FDR <0.05 and |log2 fold-change (FC)| >0.5. Top 3,000 DEGs in each subgroup are provided in STable 8. GSEA analysis was performed using ranked genes by FC, GSEA function in the clusterProfiler package^55^ (minGSSize = 20, pAdjustMethod = “BH”), and curated geneset associated with hematopoietic cells and AML generated using Human Molecular Signatures Database (MSigDB)^56^ (STable 10). Gene fusions were detected using the Genomon fusion pipeline (https://github.com/Genomon-Project/fusionfusion) and filtered for known drivers of AML.

### Bulk ATAC-seq experiments

ATAC-seq experiments were performed using Fast-ATAC protocol^18,57^. Cryopreserved tumor cells were thawed, and 50,000 cells were pelleted. For cell line experiments, K562, HL-60, SKM-1, THP-1, KY821, MOLM-13, KG-1, NOMO-1, OCI-AML3, kasumi-1, and TF-1 were included. Fifty microliters of transposase mixture (comprising 25 µL of 2x TD buffer, 2.5 µL of TDE1, 0.5 µL of 1% digitonin (Sigma-Aldrich, D141), and 22 µL of nuclease-free water) (FC-121-1030, Illumina; G9441) were added to the cells. Following transposition reactions at 37°C for 30 min, transposed DNA was purified using QIAGEN MinElute Reaction Cleanup kit or Sera-Mag™ Select (cytiva) magnetic beads, and PCR-amplified using the NEB NEXT Q5 Hot Start HiFi PCR Master Mix and custom primers^58^. TapeStation 4200 (Agilent) with High Sensitivity D5000 ScreenTape was used to assess fragment size showing nucleosomal periodicity, characteristic of the ATAC-seq library^17,59^. The resulting library was sequenced using NovaSeq 6000 instrument (Illumina) with a paired-end protocol.

### Bulk ATAC-seq analysis

Reads were aligned to the human hg19 reference genome using bowtie2^60^ with ‘-X 2000 –no-mixed –very-sensitive’ parameters, following adapter trimming using skewer^61^. The quality of sequencing data was assessed using ataqv^62^ and ATACseqQC^63^. Samples were excluded from the analysis if any of the following criteria were met: uniquely mapped reads <6M, mapping rate <40%, reads on mitochondrial chromosome >30%, or TSS enrichment scores <2 in both 1k and 2k windows. After removing duplicates and reads on the mitochondrial genome or blacklisted regions (ENCODE), peaks were called using HMMRATAC^64^ with the ‘--window 2500000’ parameter and peaks were fixed to a width of 501 bp centered on the peak summit^59^. Thereafter, unless otherwise mentioned, when peaks are extended or merged and there is overlap, the peak with the highest peak score is retained. For each of the six datasets—including our own datasets (Swedish AML, Japanese AML, AML cell lines, and normal bone marrow) as well as public datasets^18^ (sorted normal blood and AML cells)—we merged peaks from all samples within each dataset. Peaks that overlapped in two or more samples were considered the recurrent peak set for each dataset. For AML, Swedish and Japanese peak sets were further merged and filtered for those recurrently identified in both the Swedish and Japanese samples. We then merged five peak sets generated as above (AML, AML cell lines, normal bone marrow, public normal blood, and public AML) to generate a union peak set fixed to a width of 501bp, which was used for downstream analysis (ED_Fig. 1b). Annotation of peaks was conducted using the annotatePeaks function in HOMER^65^. Reads on peaks were counted and normalized to calculate CPM values using edgeR^53^. Differentially expressed ATAC peaks (DEPs) for each subgroup were identified using the eBayes test in limma through a one-versus-rest comparison, with thresholds of FDR <0.05 and |log2 FC| >0.5. Top 3,000 DEPs in each subgroup were provided in STable 9. Inference of cellular contribution to each ATAC-seq data was evaluated by the CIBERSORTx^21^ with CPM values of our AML data and public data for 13 normal blood cell types^18^ as input files.

### Clustering analysis using bulk ATAC-seq

ATAC log2 CPM were quantile normalized using preprocessCore (https://github.com/bmbolstad/preprocessCore), followed by the exclusion of peaks on chromosomes X and Y. To correct batch effects between Swedish and Japanese cohorts, ‘removeBatchEffect’ command from the limma package was used. Clustering was performed across AML samples using normalized ATAC signals on top 3,000 most variable ATAC peaks to identify ATAC subgroups. The variances of peaks across AML samples were calculated for samples with bone marrow blasts of more than 75%. The first 50 principal components were determined by prcomp in R, nearest-neighbor graph was generated using buildSNNGraph in the scran package^66^ with a ‘k = 7’ parameter, and Leiden clustering^67^ was performed using ‘cluster_leiden’ of the igraph package with parameters of ‘resolution = 0.2’ and ‘n_iterations = 100’. Clusters with less than 1% of samples (less than 16 out of 1,563 cases) were reassigned to the cluster of the nearest sample by calculating the squared Euclidean distance between samples in the two-dimensional UMAP space.

### Bulk RNA-seq-based prediction of ATAC subgroups

Normalized gene count data for four external adult AML cohorts were obtained from a previously published paper^10^. The TARGET cohort was excluded from the analysis as it was predominantly composed of pediatric AML. The count matrix was quantile normalized, and merged with our RNA-seq count matrix, followed by batch correction using ‘removeBatchEffect’. Prediction models for ATAC subgroups were generated using the Classification to Nearest Centroids (ClaNC) algorithm, which ranks genes by standard t-statistics and selects subgroup-specific genes^28^. Fifteen genes per cluster were selected for subgroup-specific markers and incorporated into the prediction model. To calculate the accuracy of prediction, we split our count matrix by cohorts and used the Swedish cohort as training and the Japanese cohort as validation. The final model to predict the ATAC subgroup in the external datasets was built using a whole cohort data.

### DNA methylation analysis

DNA methylation data (β-values) for 191 samples from the BeatAML cohort were obtained from GSE159907^15^. ATAC subgroups were predicted based on bulk RNA-seq, as described above. For heatmap generation, ATAC peak regions overlapping with methylation probes were analyzed. DNA methylation levels in a given region were calculated as the average β-values of all probes within the region. For each ATAC subgroup, average DNA methylation levels were computed, and the top 3,000 regions with the most variable DNA methylation levels across subgroups were visualized in heatmaps.

### GRN analysis by bulk ATAC– and RNA-seq

GRNs were generated using the ANANSE software^34^, utilizing the consensus ATAC peak set, merged ATAC-seq BAM file, mean RNA expression (log2 CPM), and the default motif database (GimmeMotifs^68^). GRNs were generated by calculating the number of genes each TF regulated (outdegrees) and the strength of expressional regulation (linkscores) between two TFs, by imputing 1) genome-wide binding of each TF, 2) the binding of each TF to the target genes, and 3) expression levels of each TF and its target genes (ED_Fig. 6a). GRNs were constructed for each ATAC subgroup as well as for all AML samples. The GRN of each subgroup of interest was compared to that of all AML to generate a differential (subgroup-specific) GRN and to calculate TF influence scores, which represent how well the expression differences between the two groups can be explained by a single TF (ED_Fig. 6a). For each subgroup-specific GRN, the top 20 differential TFs were drawn in the figures.

### ChIP-seq experiments and analysis

ChIP-seq experiments were performed according to the SimpleChIP Plus Sonication Chromatin IP Kit (Cell Signaling Technology)^57^ with minor modifications. Cryopreserved cells were thawed, and more than 1 million cells were fixed with 1% formaldehyde in PBS (Thermo Fisher Scientific) for 10 minutes at room temperature with gentle mixing. The reaction was stopped by adding glycine solution (10x) (Cell Signaling Technology) and incubated for 5 minutes at room temperature, and the cells were washed in cold PBS, twice. The cells were then processed with SimpleChIP Plus Sonication Chromatin IP Kit (Cell Signaling Technology) and Covaris E220 (Covaris) according to the manufacturer’s protocol. The antibodies used for ChIP were as follows: SMC1 (Abcam, ab9262), CTCF (Cell Signaling Technology, D31H2), RPB1 (CST, D8L4Y), H3K27ac (Cell Signaling Technology, D5E4), and H3K27me3 (Cell Signaling Technology, C36B11). After purification of the precipitated DNA, libraries were constructed using ThruPLEX DNA-seq kit (Takara) as per the manufacturer’s protocol, and subjected to sequencing using NovaSeq 6000 instrument (Illumina). ChIP-seq experiments were performed with input controls. The sequencing reads were aligned to the reference human hg19 genome using bowtie^69^ following adapter trimming with Skewer^61^ and read tail trimming to a total length of 50 bp using Cutadapt^70^. The quality of sequencing data was assessed using ‘plotFingerprint’ in deeptools^71^. Samples were excluded from the analysis if any of the following criteria were met: ‘X-intercept’ (deeptools) >0.85 or ‘Synthetic JS Distance’ (deeptools) <0.225. After removing duplicates and reads on blacklisted regions (ENCODE)^72^, peaks were called using MACS2^73^ and a *P*-value threshold of 1 x 10^−3^ with an input control for each sample. For each ChIP-seq (SMC1, CTCF, RPB1, H3K27ac, and H3K27me3), peaks were merged for all AML samples, and recurrently identified peaks were regarded as a consensus peak set.

### SE analysis

To identify SEs, recurrent enhancers were first identified in all AML samples using H3K27ac ChIP-seq data. Identified enhancers were stitched and ranked with H3K27ac ChIP-seq and input data, using ROSE^32^ with a ‘-t 2500’ parameter. Mean signals were used to calculate enhancer ranks across all AML samples. To identify SEs for each ATAC subgroup, mean signals were calculated to identify the top 750 enhancers in each ATAC subgroup, which were regarded as SEs. SEs were separated into three categories based on the distribution across the subgroups: Common SEs (present in 12 or more subgroups), partially shared SEs (present in 2 to 11 subgroups), and unique SEs (specific to a single subgroup). Subgroup-specific SEs were defined as those found in each subgroup and categorized as partially shared or unique SEs. Known SEs for various cell types and cancers were obtained from previous reports^32^. Annotation of SEs was conducted using annotatePeaks in HOMER^65^, filtering for genes expressed in our AML cohort (logCPM >1 in >1% of patients). AML driver genes and TF genes were determined using databases, including The Catalogue of Somatic Mutations in Cancer (COSMIC) Cancer Gene Census (CGC) (as of Nov 6th 2024)^74^, dbLGL^75^, TF list from a previous study^76^, and the Jaspar^18,77^ database, and were manually selected. A list of SEs identified in this study is summarized in STable 6.

### SE-regulated gene signature

Enriched gene ontologies in SE-associated genes were identified using enricher in the clusterProfiler package^55^ (pAdjustMethod = “BH”, qvalueCutoff = 0.25) and ontology geneset (‘hallmark’, ‘c2.cp.reactome’, and ‘c5.go.bp’) from MSigDB (v2024.1)^56^. Expression levels of SE-associated genes in each hematopoietic cell type were calculated using public gene expression data from DMAP^44^, and the average expression levels were determined for each cell type.

### SE-based TF network analysis

To analyze SE-regulated TF networks, we applied the coltron algorithm^35^ with ROSE^32^ outputs generated from the merged BAM files for each ATAC subgroup. This algorithm computes the inward binding (in-degree) of other SE-associated TFs to a given SE-associated TF, as well as the outward binding (out-degree) of the TF to other SEs. The coltron score for each TF was determined as the sum of its in-degree and out-degree (ED_Fig. 6c).

### scRNA/ATAC-seq experiment

Single-cell matched RNA-seq and ATAC-seq experiments were performed using the Next GEM Single Cell Multiome ATAC + Gene Expression Reagent Kit (10x Genomics), according to the manufacturer’s protocols (CG000365 for nuclei isolation and CG000338 for library generation). Cryopreserved cells were thawed, dead cells were stained with DAPI, and live MNCs were sorted using FACS Aria III (BD Biosciences). The samples were resuspended in lysis buffer (10 mM Tris-HCl (pH 7.4), 10 mM NaCl, 3 mM MgCl2, 1% BSA (Miltenyi Biotec,130-091-376), 0.1% Tween-20 (Bio-Rad, 1610781), 0.1% IGEPAL (Sigma-Aldrich, i8896), 0.01% digitonin (Thermo Fisher, BN2006), 1 mM DTT (Sigma-Aldrich, 646563), plus 1 U/μl RNase Inhibitor (ThermoFisher, 10777019)), incubated on ice for 3 minutes, washed with wash buffer (10 mM Tris-HCl (pH 7.4), 10 mM NaCl, 3 mM MgCl2, 1% BSA, 0.1% Tween-20, 1 mM DTT, plus 1 U/ μl RNase Inhibitor) three times, and passed through a 40 μm Flowmi Cell Strainer (Bel-Art). After microscopy inspection and counting of nuclei using a Countess II FL Automated Cell Counter (Thermo Fisher Scientific), nucleus suspensions were prepared in a concentration targeting a maximum of 10,000 nuclei recovery, and incubated with transposase to add adapter sequences to the DNA fragments. The suspensions containing transposed nuclei were subjected to the gel beads in emulsion (GEMs) generation, incubation, and cleanup, using the Chromium Next GEM Chip J Single Cell Kit (10x Genomics) and Chromium Controller. The resulting suspensions contained ATAC fragments and cDNA with the same cell barcodes. The pre-amplification of cDNA was performed, and the amplified product was split and used as the input for ATAC and gene expression library construction. Libraries were generated using the 10x Genomics Single Index N Set for ATAC and the 10x Genomics Dual Index TT Set A for RNA. The scATAC libraries were sequenced using DNBSEQ G400 (MGI) instrument with a custom protocol (Read 1: 50 cycles, Read 2: 49 cycles, i5 Index: 24 cycles, i7 Index: 8 cycles). The scRNA libraries were sequenced using DNBSEQ G400 (MGI) instrument with a custom protocol (Read 1: 28 cycles, Read 2: 90 cycles).

### scRNA/ATAC-seq analysis

Cell Ranger ARC (10x Genomics)^78^ was used for data processing to generate bam files and count matrices for scRNA and scATAC with the hg19 reference genome and GENCODE human v19 annotation^79^. The report generated by Cell Ranger was manually evaluated and samples were filtered to remain with no errors or with only warnings. Basic QC reports including analyzed cell numbers from Cell Ranger for each sample were summarized in STable 7. ArchR^80^ was used to filter for cells with following thresholds: number of unique Molecular Identifiers >100 and proportion of mitochondria genes <0.05 for scRNA; transcription start site enrichment score >4 and number of fragments >1000 for scATAC. Doublet cells were removed by the ‘filterDoublets’ function in ArchR with a filterRatio of 1. The gene expression count matrix was stored in the Seurat^81^ object, and mitochondrial genes were excluded from the following analysis. The consensus peak set, generated from bulk ATAC-seq as previously mentioned, was filtered for peaks on autosomal chromosomes and used to count reads on peaks in the scATAC analysis, utilizing the FeatureMatrix function from the Signac package^82^.

### Merging individual data, normalization, and clustering for scRNA/ATAC-seq

Raw count matrix data for single cell gene expression and chromatin accessibility was written to disk using the BPCells package (https://github.com/bnprks/BPCells) to enable high-throughput data processing and merged across samples. The merged gene count matrix was normalized for sequencing depth using the SCTransform function from Seurat^81^. The merged count matrix for scATAC was normalized using the term frequency inverse document frequency (TF-IDF) normalization function in BPCells. Cells were then clustered based on scATAC using the top 10 PCA dimensions with knn_hnsw (ef=1000), knn_to_snn_graph, and cluster_graph_louvain (resolution=0.1) functions in BPCells. Each scATAC cluster was classified as the AML predominant cluster if they met any of the following criteria: 1) >80% of cells in the cluster were derived from AML samples, or 2) >50% of cells in the cluster were derived from AML samples and >30% of cells in any single AML sample were assigned to the cluster. Additionally, each cluster was excluded from AML clusters and regarded as the normal predominant cluster if they met either of the following criteria: 1) >10% of cells in each CR sample were assigned to the cluster, or 2) <30% of cells in each AML subgroup were assigned to the cluster.

### Estimation of differentiation and pseudotime using scRNA-seq

The BoneMarrowMap package in R was used to project AML cells onto reference scRNA-seq data of human bone marrow hematopoiesis^40^. Cells with low mapping quality were excluded from the analysis based on the following criteria: 1) Mean absolute deviation of mapping error scores ≧2; 2) Assigned to ‘Orthochromatic Erythroblast’ and module score of hemoglobin genes <1.5. The module score was calculated using the ‘AddModuleScore’ function in Seurat^81^ with hemoglobin genes. Pseudotime analysis was subsequently performed using the ‘predict_Pseudotime’ function in BoneMarrowMap.

### Single-cell GRN analysis

The SCENIC+ software^41^ (v1.0a1) was used to evaluate TF activities with combined scRNA-seq and scATAC-seq data, following the standard vignettes with minor modifications (ED_Fig. 8a). Cells were filtered for AML samples and those categorized in scATAC AML clusters and used for making eRegulons. One hundred topics were empirically selected. The pipeline was run in multiome mode, using 5 cells per metacell. The search space was defined as 0-500kb. Established eRegulons were filtered with the following parameters: ‘rho_threshold’=0.03, ‘min_regions_per_gene’=0, and ‘min_target_genes’=10. To calculate region-based and gene-based specificity scores in ATAC subgroups, cells were downsampled to 20,000 in each subgroup. To calculate scores in each pseudotime bin in each ATAC subgroup, cells were downsampled to 300 if the number of cells was more than 300 in each bin. eRegulons with positive ‘TF-to-gene’ and ‘region-to-gene’ relationships were used for downstream analysis. Only direct eRegulons were kept and extended ones were added only if there was not a direct one.

### AML patient sample drug screening

The procedure for drug sensitivity and resistance testing in the samples used for the study has previously been described in detail^43^. Biobanked MNCs were thawed and added to pre-spotted drug plates (FIMM HTB)^83^. The MNCs were incubated in HS-5 conditioned (12.5%, ATCC, Manassas, VA) complete RPMI (10% FBS (Thermo Fisher), 2 mM L-glutamine (Sigma-Aldrich), 100 IU/mL Penicillin and 0.1 mg/mL Streptomycin (Pen-Strep, Sigma-Aldrich), for 72 h (37 °C, 5% CO_2_). Cell viability was measured by CellTiterGlo (CTG, Promega, Madison, WI) on an EnSight plate reader (PerkinElmer, Waltham, MA). Drug sensitivity scores (DSS) were calculated with Breeze^84^. Selective DSS (sDSS) values were calculated by subtracting healthy BM control DSS values from patient DSS values.

### Statistical analysis

Statistical analyses were performed using the R software. Comparisons between groups were based on the two-sided Wilcoxon rank-sum test for continuous data and the Fisher’s exact test for categorical data, unless otherwise specified. Survival analysis was performed for patients who were treated with intensive chemotherapy, and observations were censored at the last follow-up. The Kaplan-Meier method was used to estimate the OS, and the differences in OS were assessed by the log-rank test. The effects of clinical variables and ATAC subgroups on OS were evaluated using the Cox proportional hazards regression model for each ELN risk. For clinical parameters, the median values were used as the threshold unless otherwise specified. The prognostic model was established with the training (Swedish) cohort, and the performance of the model was validated by calculating the concordance statistic (c-index) with bootstrapping 1,000 times in both training and validation (Japanese) cohorts. Explained variances in several features by ICC, WHO, and ATAC classifications were evaluated as *r^2^* values calculated by the adonis function in R, with Euclidean distances. For gene expression profiles, the first 50 principal components were used as input.

## Data and code availability

The datasets generated in this study will be available upon reasonable request and will be deposited in a public repository upon publication. ATAC-seq data for sorted normal blood and AML cells were obtained from Gene Expression Omnibus (GEO) database under the accession number GSE74912^18^. DNA methylation data for BeatAML was obtained from GSE159907^15^. Normalized gene count data based on RNA-seq used for prediction of subgroups were obtained from the previous paper^10^. Public dataset used in this study is summarized in STable 5.

## Acknowledgments

We are grateful to all the patients and their families for their contribution of biological specimens to this study. We also appreciate the clinicians and healthcare professionals who provided biospecimens and contributed to the biobanks in Sweden and Japan. Additionally, we acknowledge N. Sakamoto, T. Shirahari, S. Yabuta, H. Takada, K. Onishi, and R. Onoi, as well as the Single-cell Genome Information Analysis Core (SignAC) at WPI-ASHBi, Kyoto University, for their technical support.

This work was supported by grants from the Japan Science and Technology Agency (JST): FOREST Program (no. JPMJFR220L) to Y.O.; the Japan Agency for Medical Research and Development (AMED): nos. JP21cm0106501, JP19ck0106250, and JP24tk0124003 to S.O, JP24ck0106791 and JP25ck0106019 to Y.O. and H.I.S, JP25kk0305028 to Y.O., JP22ck0106691 to Y.O., N.Y., and S.O, JP22ama221111 and JP23kk0305026 to H.I.S., JP25ama121016 to H.A., and JP24ck0106875 to Y.O., N.Y., H.I.S., and S.O; the Moonshot Research and Development Program (no. JP24zf0127009) to S.O.; the Ministry of Education, Culture, Sports, Science and Technology of Japan (MEXT): nos. hp200138, hp210167, JPMXP1020200102 to S.O. and M.S; the Japan Society for the Promotion of Science (JSPS): KAKENHI (nos. JP19H05656, JP24H00009 to S.O, JP20K22809, JP22K16320, JP24K19223 to Y.O.); Takeda Science Foundation to S.O. and Y.O.; The Chemo-Sero-Therapeutic Research Institute, Daiichi Sankyo Foundation of Life Science, Princess Takamatsu Cancer Research Fund, Kanae Foundation for the Promotion of Medical Science, Ichiro Kanehara Foundation for the promotion of Medical Sciences and Medical Care, Mochida Memorial Foundation for Medical and Pharmaceutical Research, Japan Leukemia Research Fund, Kobayashi Foundation for Cancer Research, Fujiwara Memorial Foundation, and UBE Foundation to Y.O.

## Author contributions

Y.O., S.L., and S.O. designed the study. M.L.-L., S.B., B.P., S.D., M.J., V.L.,J.C., A.R., L.W., E.O., S. Kasahara, N.H., N.K., N. Sezaki, M.S., M.I., J.K., Y.U., S.Y., T.I., S. Matsuda, A.T.-K., G.J., and M.H. provided patient samples and collected clinical data. Y.O. and Y.N. performed targeted-capture sequencing. Y.O. performed RNA-seq, ATAC-seq, ChIP-seq, and scRNA/ATAC-seq. S. Morimoto and H.A. performed DNA methylation array. T.E., N. Struyf, and O.K. performed drug screening. Y.O., Y.N., M.M.N., R.S., H.F., K.O., A.Y., R.O., G.X., A.O., A.K., L.Z., Y.S., and S. Miyano analyzed genomic data. Y.O., Y.W., M.M., S, Komatsu, and H.I.S. analyzed epigenetic data. Y.O. and S.O. wrote the manuscript. S.O. and S.L. jointly lead the study. All authors reviewed and approved the manuscript.

---

**Extended Data Fig. 1.**
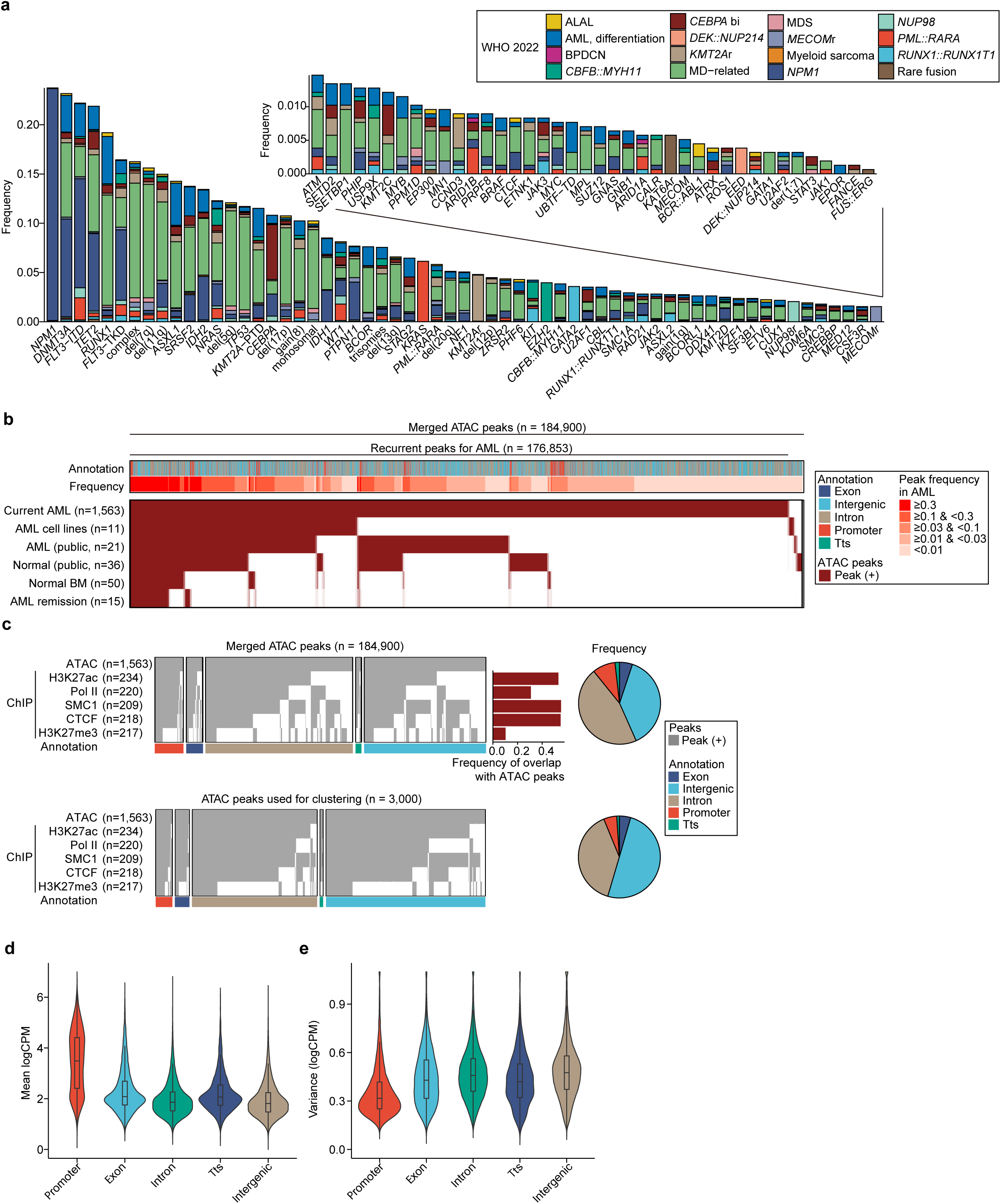
Landscape of genome and epigenome in AML. **a.** Frequencies of driver mutations in 1,563 AML patients. Each color represents WHO 2022 diagnosis. **b.** Summary of ATAC peaks recurrently identified in each dataset with annotations and frequencies in AML samples. **c.** Summary of all ATAC peaks (top panel) and those used for ATAC-based clustering (bottom panel), with annotations and overlaps with ChIP-seq peaks. Pie charts show proportions of each annotation. **d-e.** Mean (**d**) and variance (**e**) of ATAC signals in each peak type across all AML patients.

**Extended Data Fig. 2.**
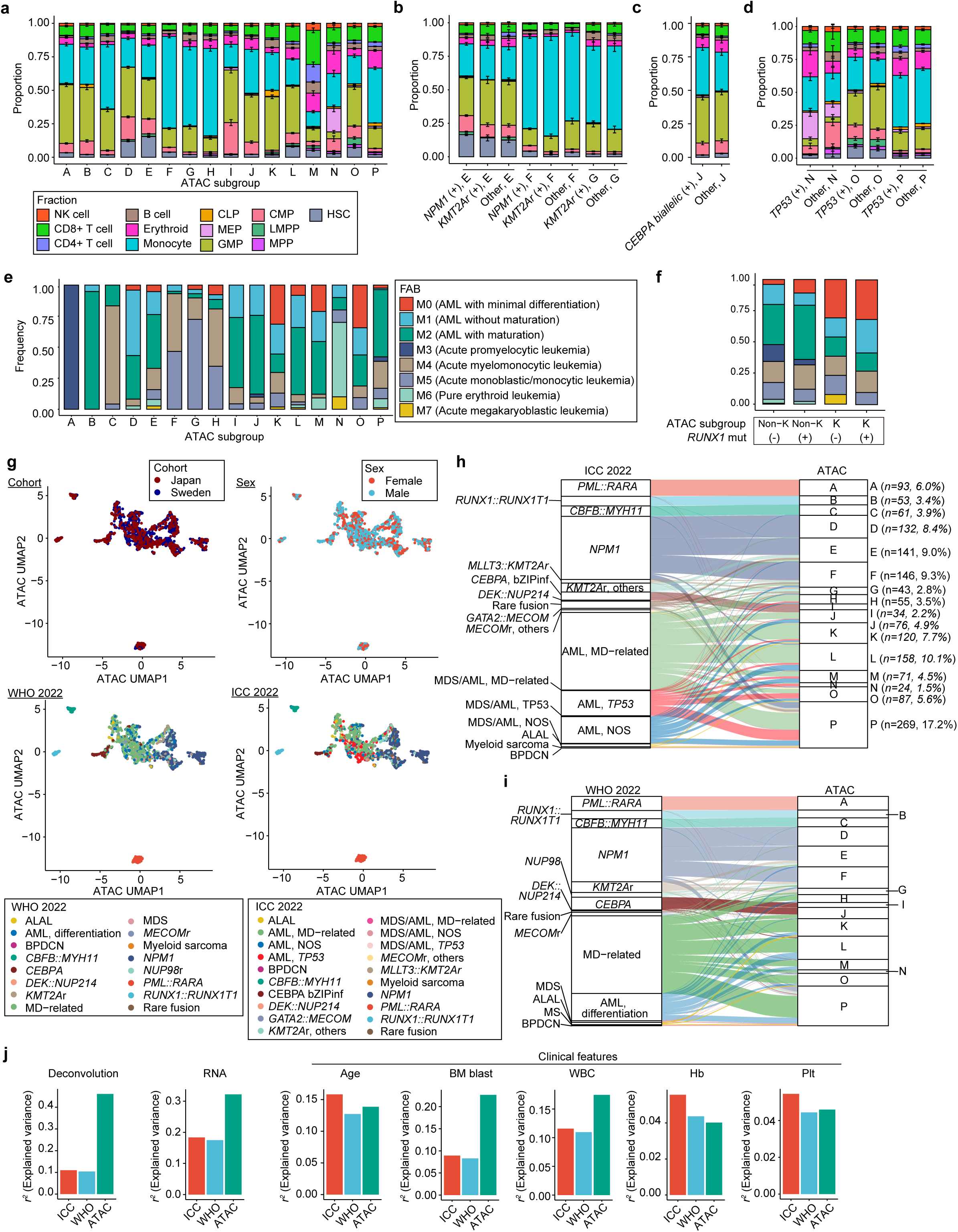
Epigenetic subgroups and differentiation states. a-d. Contributions from each blood cell type in each bulk ATAC-seq data according to ATAC subgroups, estimated by CIBERSORT. ATAC subgroups are further separated by the presence of *NPM1* mutation or *KMT2A* rearrangement (**b**), *CEBPA* biallelic mutations (**c**), and *TP53* mutations (**d**). **e-f.** Proportion of each FAB diagnosis in ATAC subgroups (**e**) and in subgroup K with or without *RUNX1* mutations (**f**). **g.** UMAP plot based on ATAC-seq profiles. Each dot represents one patient and colors indicate cohort (top left), sex (top right), WHO 2022 (bottom left), and ICC 2022 (bottom right) diagnoses. **h-i.** Sankey plot showing the assignment of ICC 2022 (**h**) and WHO 2022 (**i**) diagnoses over the identified ATAC subgroups. **j.** Bar plots showing the explained variances (*r*_2_ values calculated by permutational multivariate analysis of variance) for each feature by WHO, ICC, and ATAC classifications.

**Extended Data Fig. 3.**
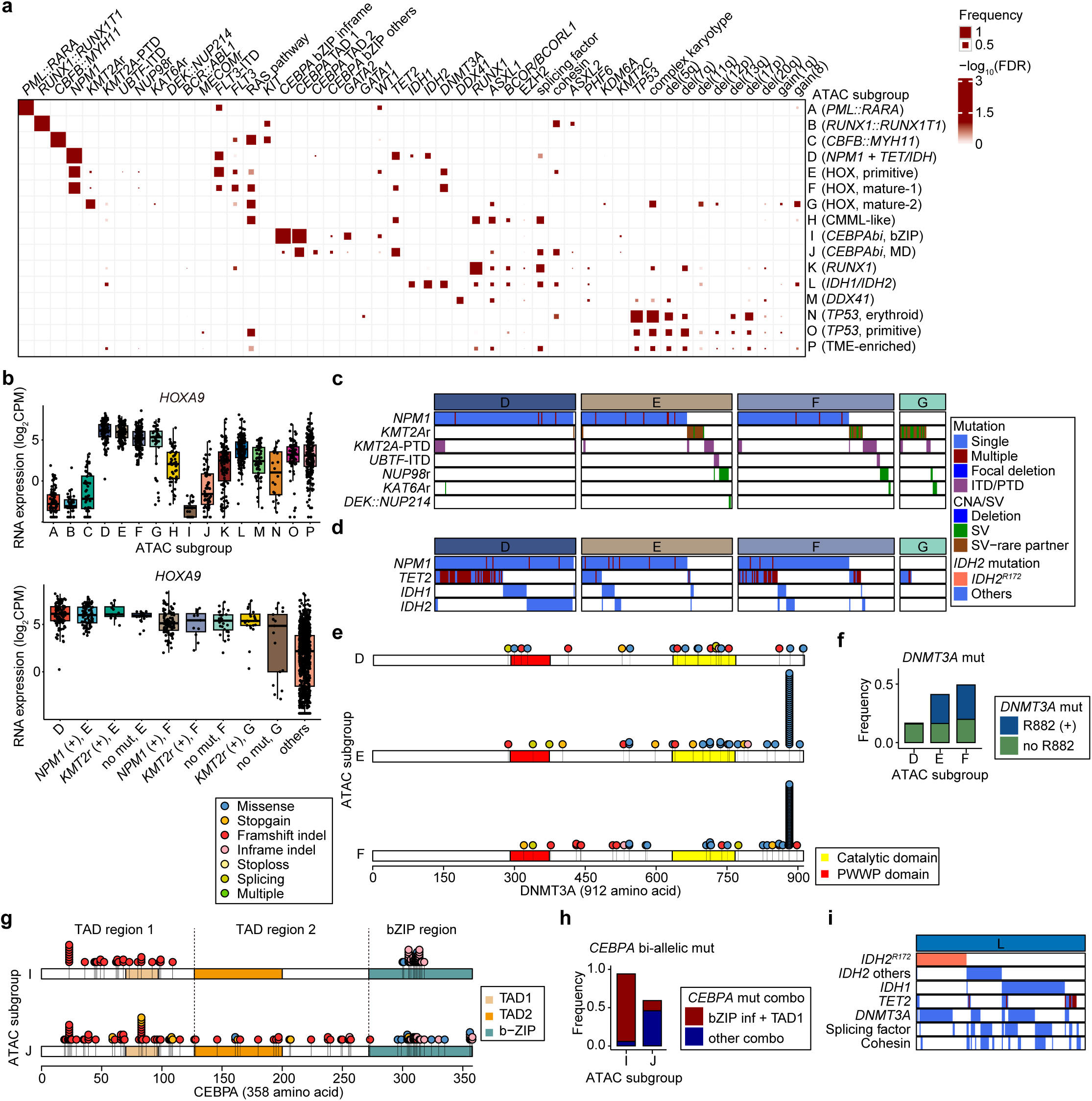
Unique co-mutations enriched for each ATAC subgroup. **a.** Frequencies of genetic abnormalities in each subgroup. *P* values for enrichment of each genetic abnormality were calculated using a one-sided Fisher’s exact test and adjusted for multiple comparisons with the Benjamini-Hochberg method (FDR). * FDR <0.05; ** FDR <0.01; *** FDR <0.001. **b.** RNA expression of *HOXA9* according to ATAC subgroups (top boxplot). ATAC subgroups were further separated by the presence of *NPM1* mutation or *KMT2Ar* (bottom boxplot). CPM, counts per million. **c,d,i.** Summary of co-occurrence of genetic abnormalities in subgroups D-G (**c,d**), and subgroup L (**i**). Column represents one patient, sorted by the presence of indicated genetic abnormalities. **e,g.** Distribution of mutations in *DNMT3A* (**e**) and *CEBPA* (**g**) genes in each ATAC subgroup. **f.** Frequency of R882X (blue) and other (green) *DNMT3A* mutations in subgroups D-F. **h.** Frequency of *CEBPA* mutation pairs in biallelic *CEBPA* mutations.

**Extended Data Fig. 4.**
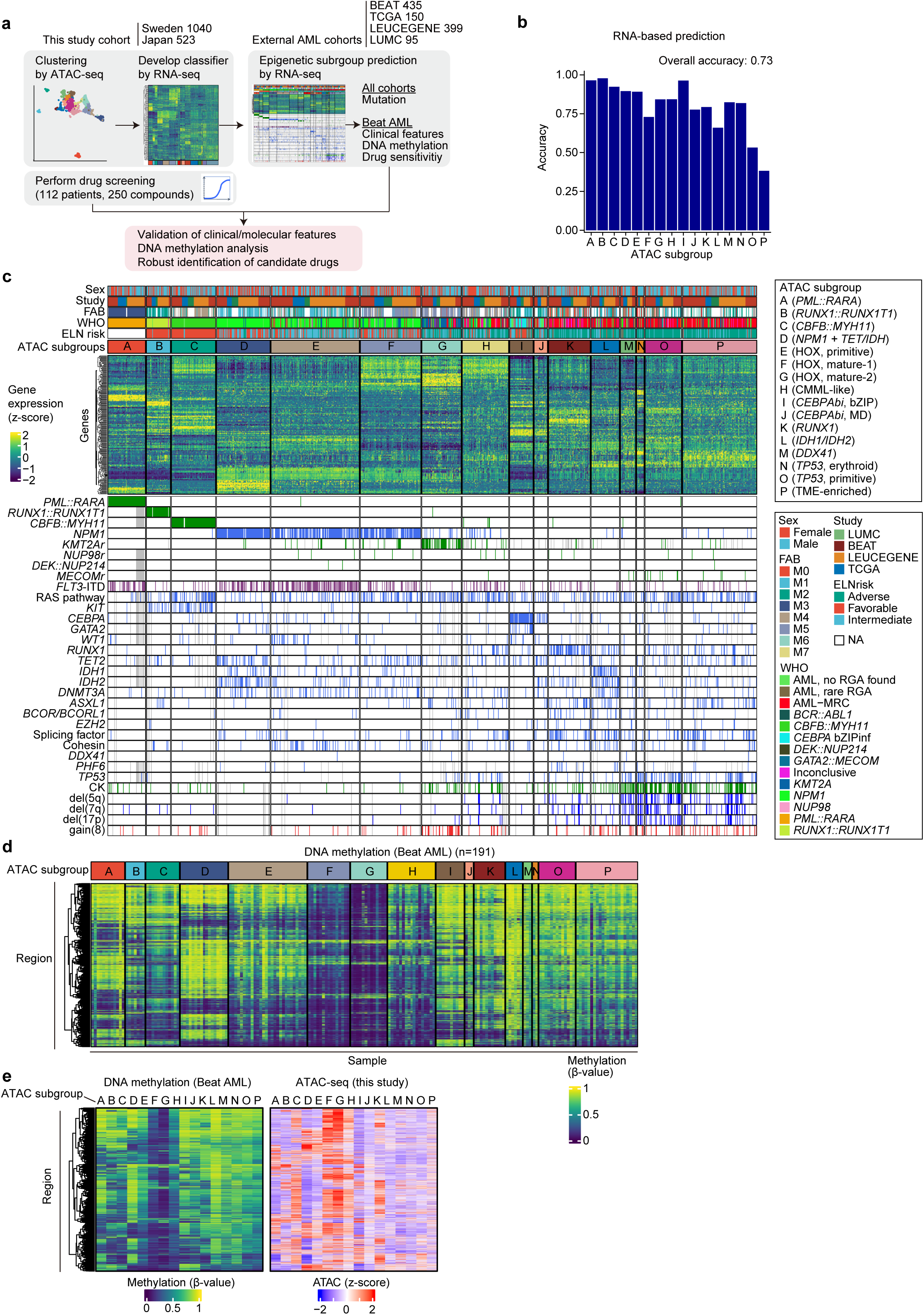
Prediction of ATAC subgroups using gene expression profiles. **a.** Scheme of ATAC subgroup prediction and downstream analysis. **b.** Prediction accuracy for each ATAC subgroup. The accuracy of prediction was calculated using the Swedish cohort as training and the Japanese cohort as validation. Overall accuracy was calculated for all samples. **c.** Summary of clinical features, diagnoses, predicted ATAC subgroups, scaled expression of marker genes that were defined by ClaNC algorithm, and genetic abnormalities for predicted ATAC subgroups for four external adult AML cohorts (LUMC, BEAT, LEUCEGENE, and TCGA). **d.** DNA methylation levels of 3,000 variable regions in each sample in the BeatAML dataset. **e.** Mean DNA methylation levels of 3,000 variable regions in each predicted ATAC subgroup in the BeatAML dataset are shown on the left. Mean ATAC signals for the corresponding regions in each ATAC subgroup from our cohort are shown on the right.

**Extended Data Fig. 5.**
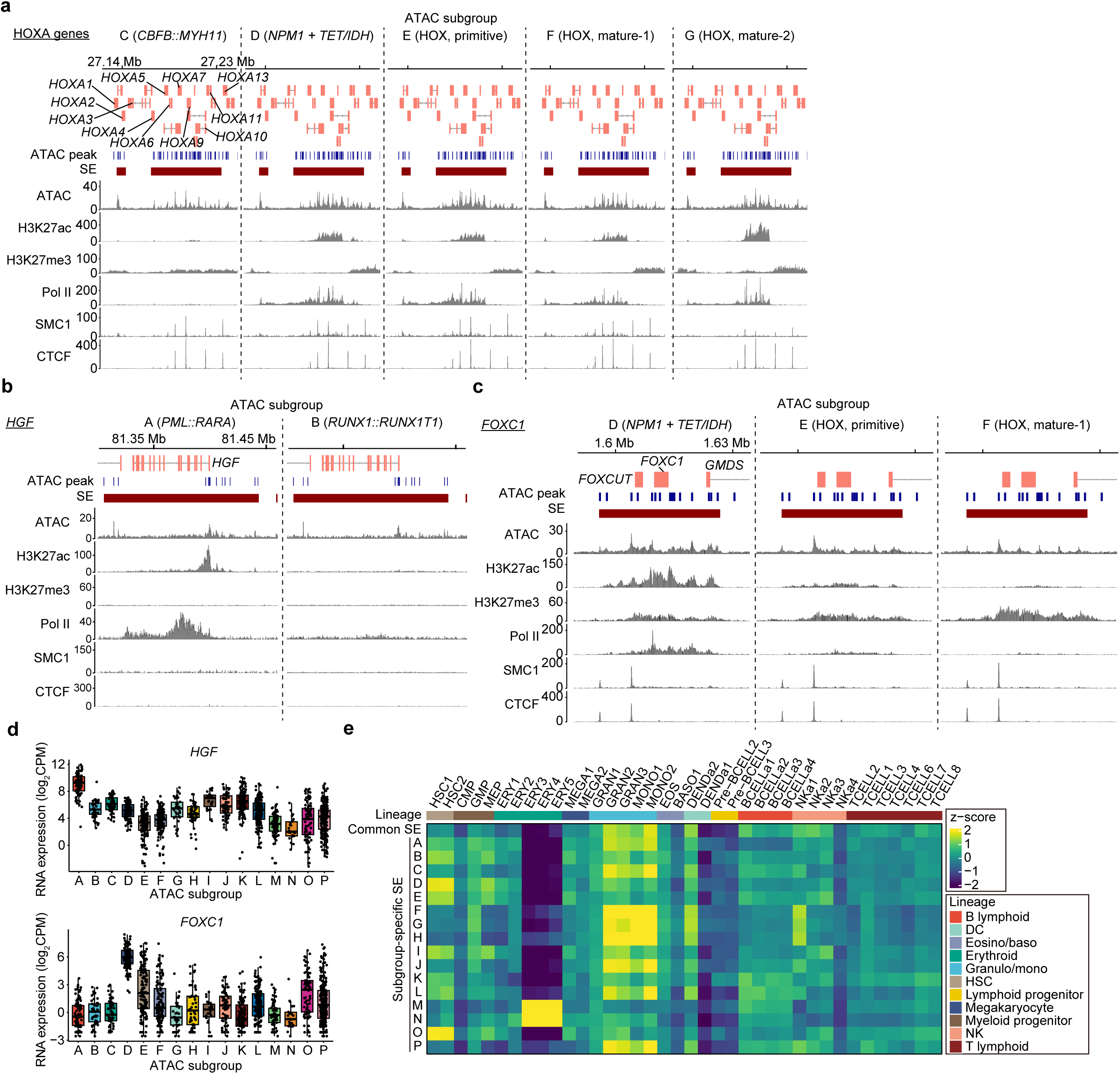
Subgroup-specific SEs and lineage specificity. a-c. ATAC-seq and ChIP-seq tracks showing the average signals for the indicated subgroups around the specified SEs. **d** RNA expression of SE-regulated genes, *HGF* (top) and *FOXC1* (bottom), according to subgroups. CPM, counts per million. **e.** Scaled expression levels of genes associated with common and subgroup-specific SEs in each hematopoietic cell type_50_. Each column represents cell type, and each row shows common SE and subgroup-specific SEs in each subgroup.

**Extended Data Fig. 6.**
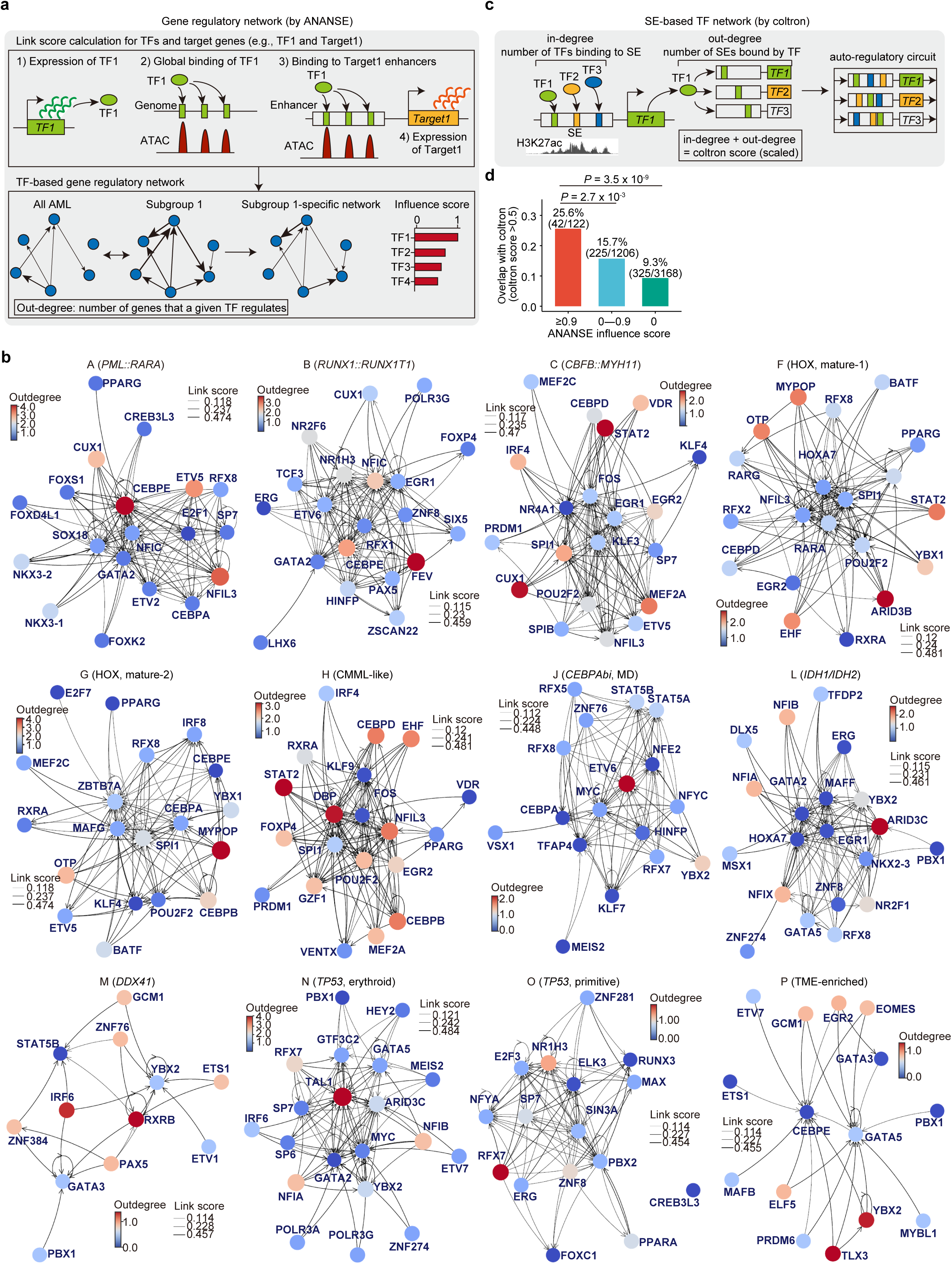
Network analysis combining multiomic epigenetic analysis. **a.** Scheme for gene regulatory network analysis using the ANANSE software_38_. **b.** GRNs specific to the indicated subgroups, generated using ANANSE. Colors indicate the regulatory strength of the corresponding TFs on other TFs (outdegree). Line width represents the degree of interaction between two TFs (link score). **c.** Scheme for SE-based TF network analysis using the coltron software_39_. **d.** Overlap ratios between the ANANSE-based network and coltron-based network. The overlap ratio was calculated as the proportion of instances where both the Coltron score (>0.5) and the ANANSE influence score (indicated range) met their respective thresholds across all combinations of subgroups and TFs. *P* values were calculated by Fisher’s exact test.

**Extended Data Fig. 7.**
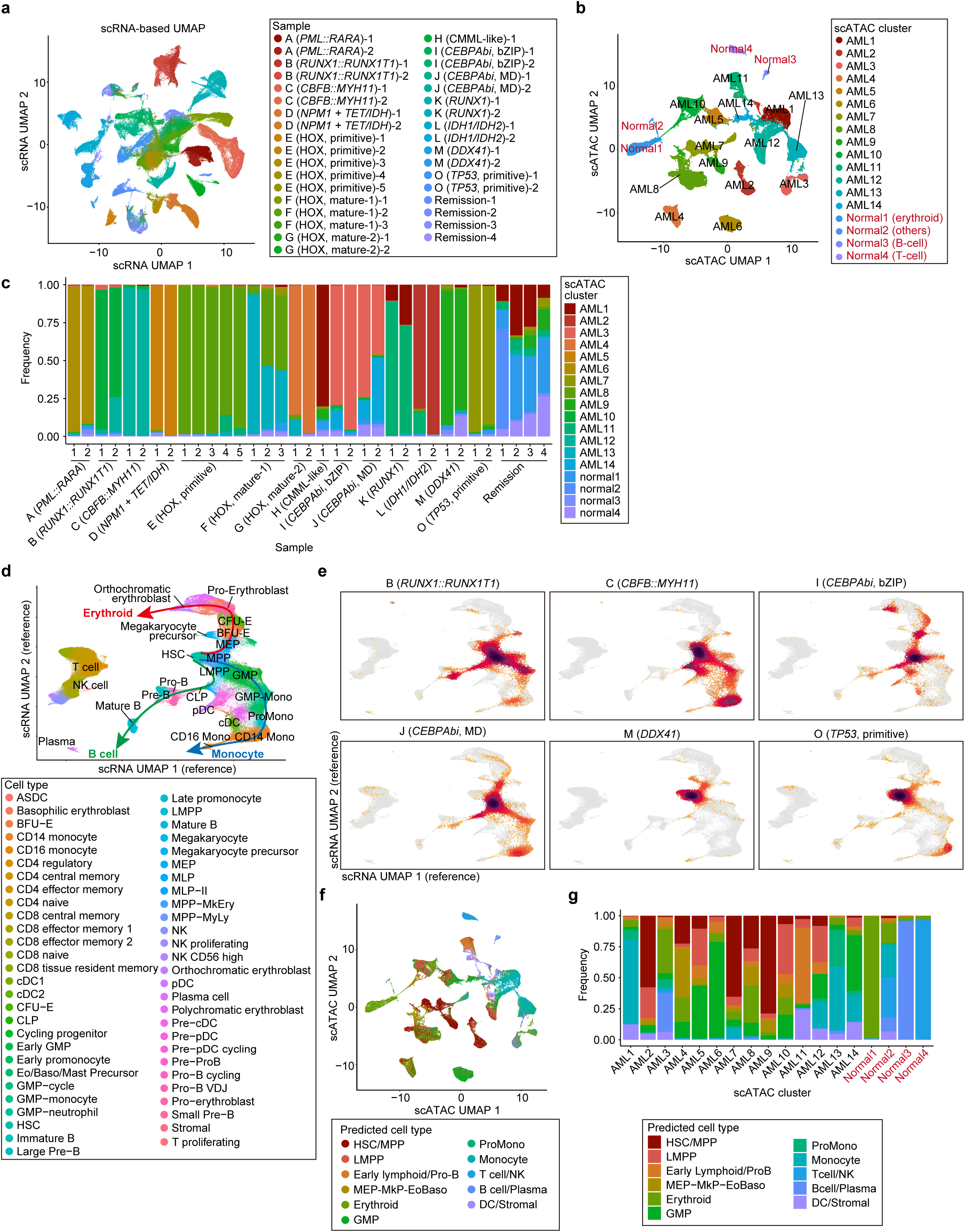
scRNA/ATAC-seq profiles and predicted differentiation status of AML cells. **a.** UMAP plot based on scRNA-seq profiles. Each dot represents one cell, colored by samples. **b.** UMAP plot based on scATAC-seq profiles. Each dot represents one cell, colored by clusters determined by scATAC-based clustering. **c.** Proportion of cells assigned to scATAC-seq clusters for each patient sample. **d.** Scheme and annotated cell types for reference-based pseudotime analysis. ASDC, AXL+ Siglec-6+ dendritic cells; BFU-E, burst forming unit erythroid; CFU-E, Colony forming unit erythroid; Eo, eosinophil; Baso, basophil; LMPP, Lymphoid-primed multipotent progenitor; MLP, multi-lymphoid progenitor; MPP, multipotent progenitor; pDC, plasmacytoid dendritic cell; cDC, conventional dendritic cell; VDJ, VDJ recombination. **e.** Mapping of AML cells from each ATAC subgroup onto the reference scRNA-seq UMAP space. **f.** UMAP plot for scATAC-seq profiles, with colors indicating predicted cell types. **g.** Proportions of estimated cell types in each scATAC cluster.

**Extended Data Fig. 8.**
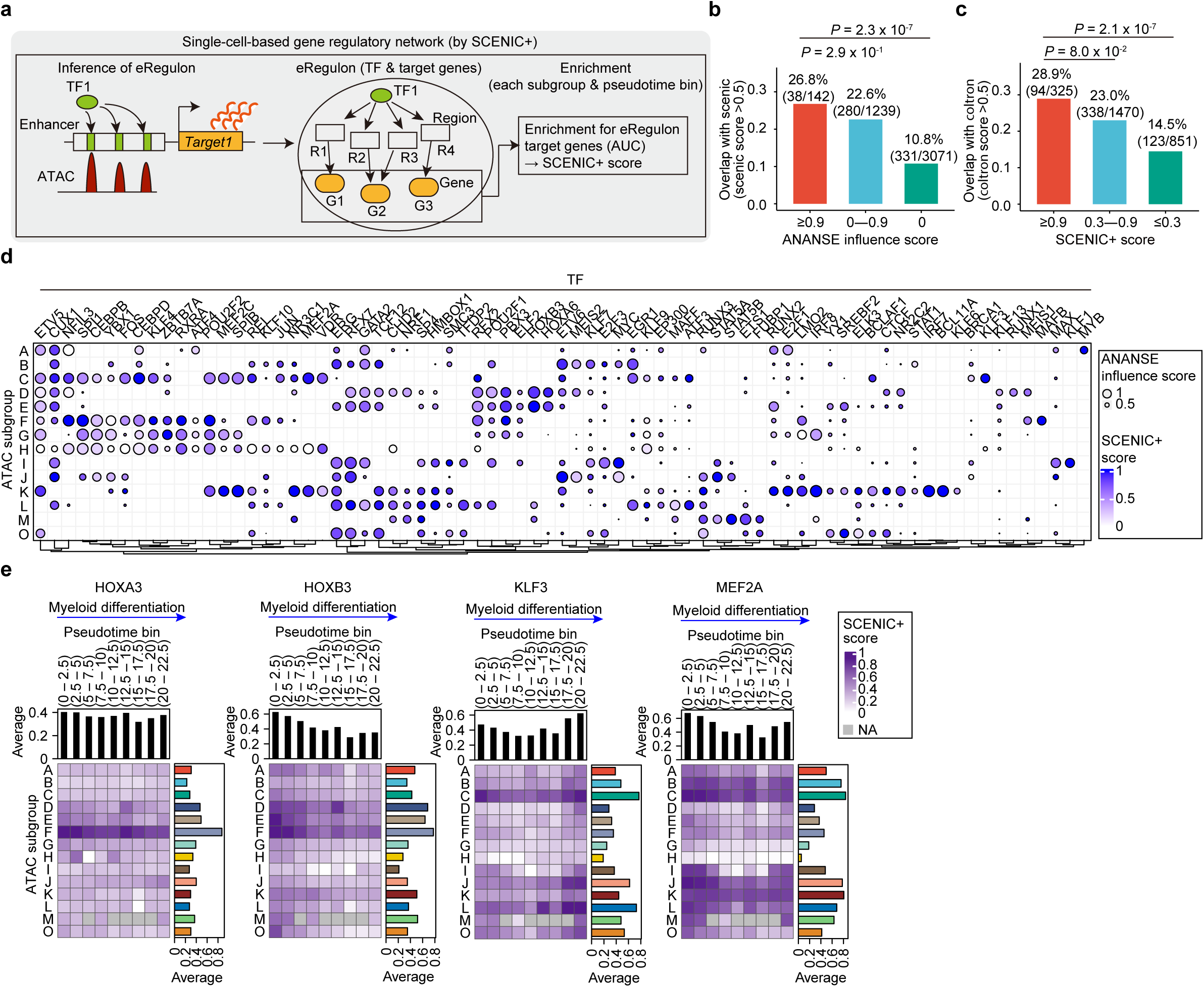
TF activities at the single-cell level. **a.** Scheme for TF activity estimation using the SCENIC+ software_47_. **b-c.** Overlap ratios between the SCENIC+-based network and ANANSE/coltron-based networks. The overlap ratio was calculated as the proportion of instances where scores from both pipelines met their respective thresholds across all combinations of subgroups and transcription factors (TFs). *P* values were calculated by Fisher’s exact test. **d.** Heatmap summarizing activities of TFs that were identified in both ANANSE and SCENIC+ analysis. Size indicates influence scores for each TF, calculated by the ANANSE software_38_ (ED_Fig. 6a). Color indicates the SCENIC+ score (gene-based Area Under the Curve (AUC) in SCENIC+ software_47_ (ED_Fig. 8a)). Each row represents the ATAC subgroup. **e.** TF activities inferred by SCENIC+ scores in each pseudotime bin along the myeloid differentiation trajectory.

**Extended Data Fig. 9.**
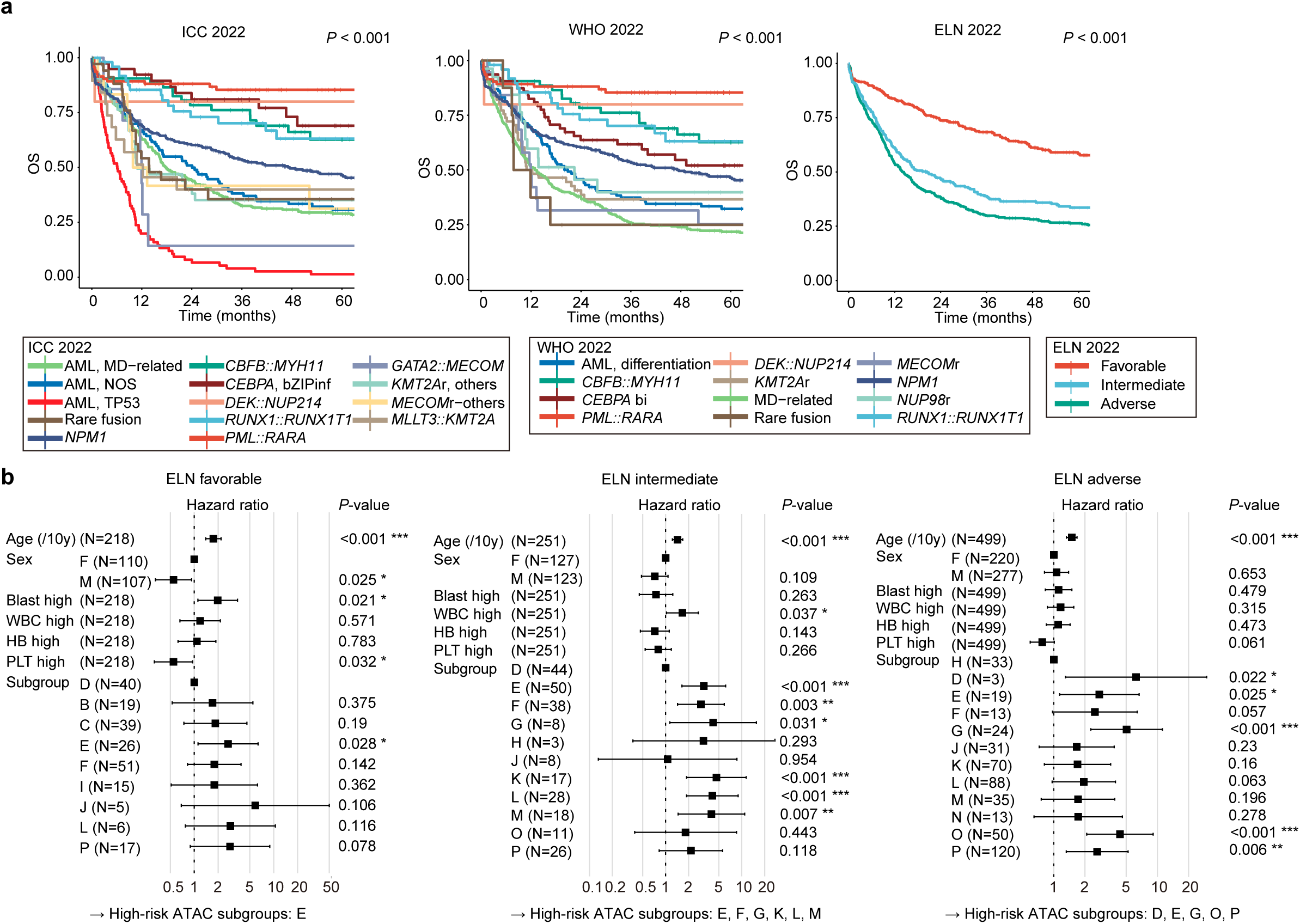
Survival analysis using combined genome and epigenome data. **a.** Kaplan–Meier survival curves for OS of patients who received intensive chemotherapy, categorized by ICC 2022 (left), WHO 2022 (center), and ELN2022 (right). *P* values were calculated using the log-rank test. **b.** Cox regression analysis for patients in each ELN risk group, incorporating clinical variables and ATAC subgroups with more than two patients. In each analysis, the subgroup with the most favorable survival was used as the reference, and high-risk ATAC subgroups were defined as those with significantly worse prognosis.

**Extended Data Fig. 10.**
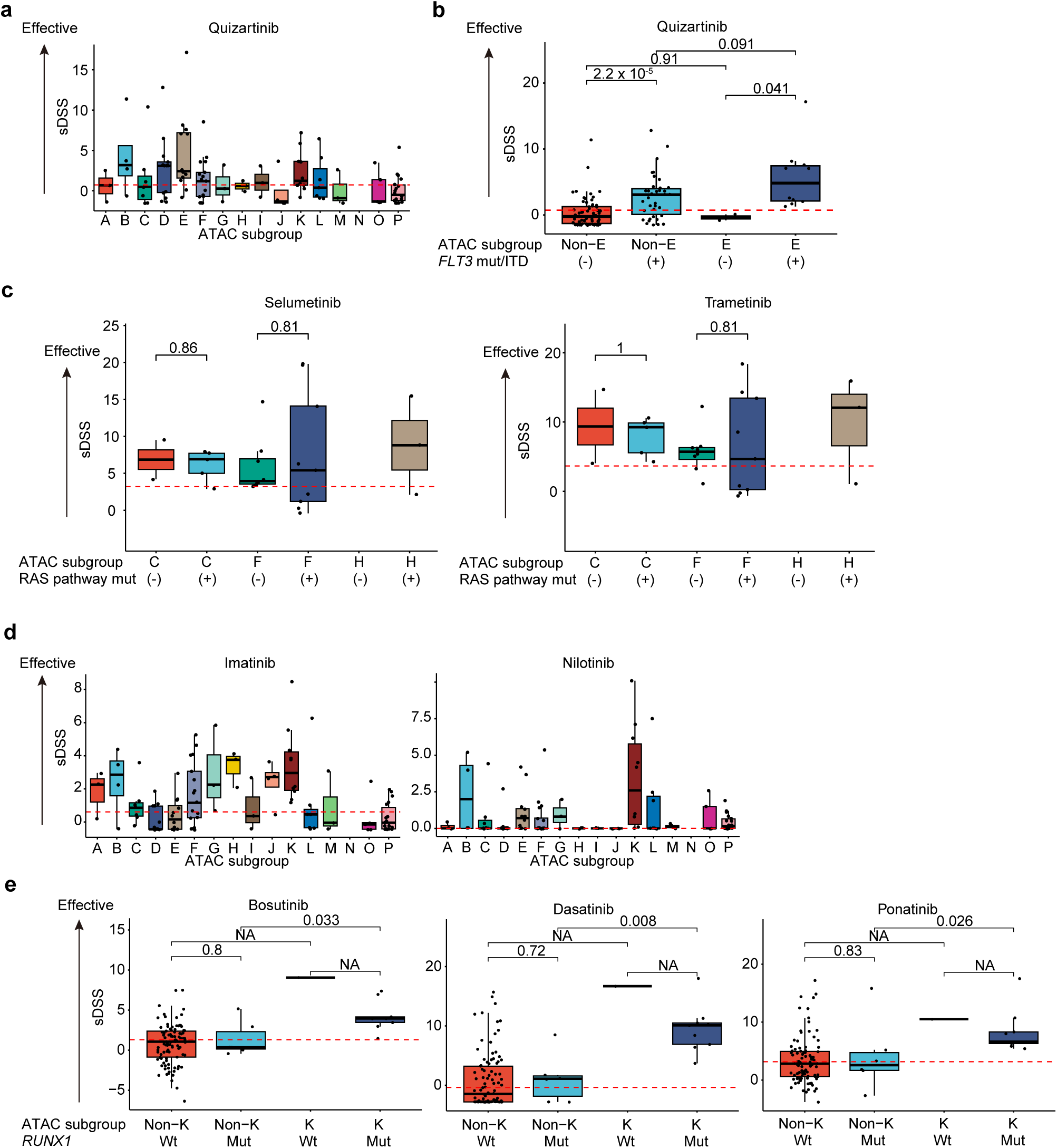
Drug sensitivities according to mutations and ATAC subgroups. a,d. Sensitivity to FLT3 (**a**) and ABL inhibitors (**d**) in each subgroup. **b,c,e.** Sensitivity to FLT3 inhibitors (**b**), MEK inhibitors (**c**), and ABL inhibitors (**e**) based on ATAC subgroups and the presence of indicated mutations. The dashed line indicates the median value across all patients. *P* values were calculated using the two-sided Wilcoxon rank-sum test. Wt, wild-type; Mut, mutated.

