## Supplementary Figures for "Chromatin landscape and epigenetic heterogeneity of acute myeloid leukemia"

#### Supplementary Fig. 1

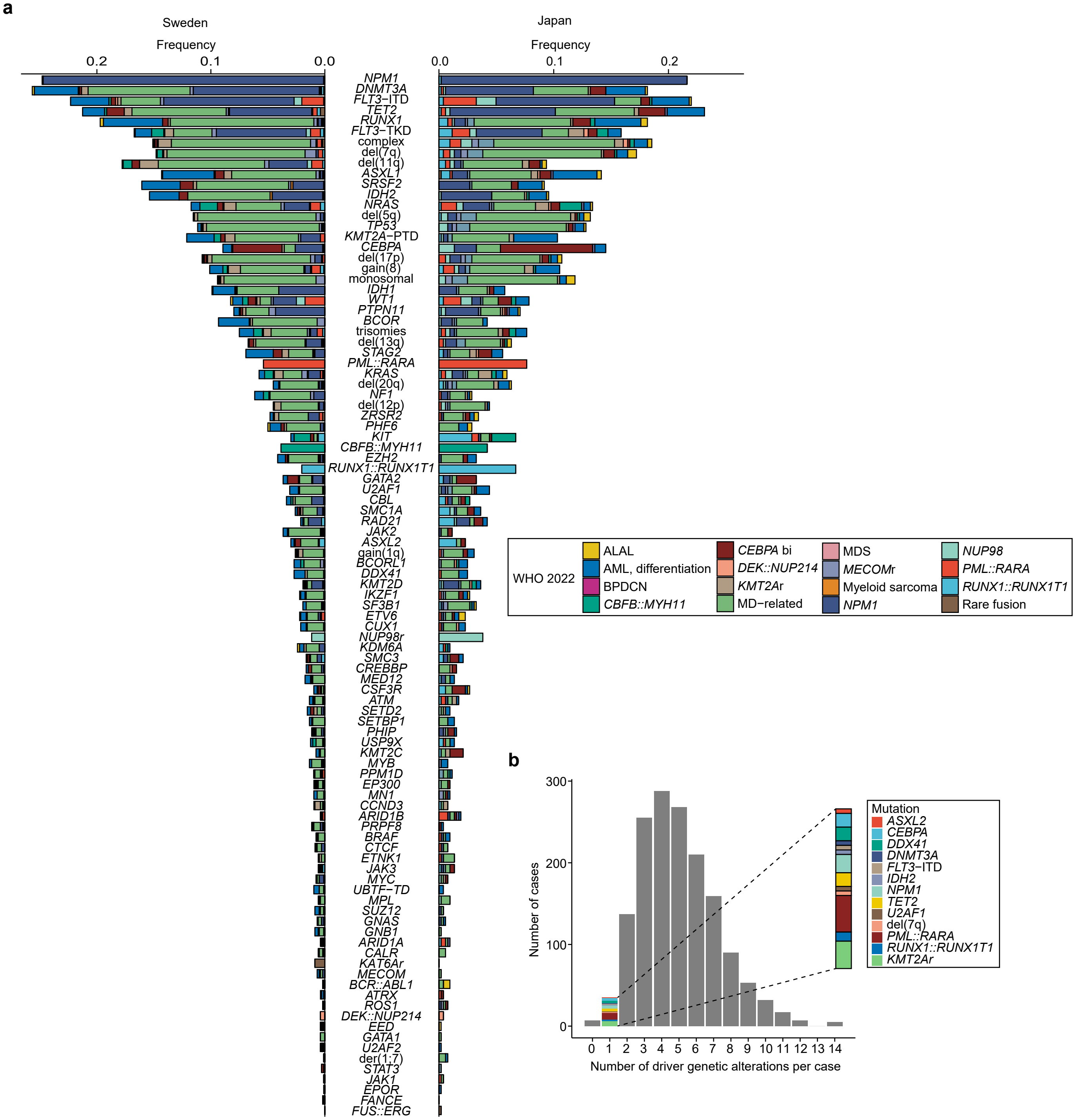


**Supplementary Fig. 1 Summary of driver genetic alterations. a.** Frequencies of genetic alterations in Swedish and Japanese cohorts. Each color represents WHO 2022 diagnosis. **b.** Number of driver genetic alterations per sample is shown as a barplot. In samples with only one alteration, the specific alteration is highlighted.

### Supplementary Fig. 2

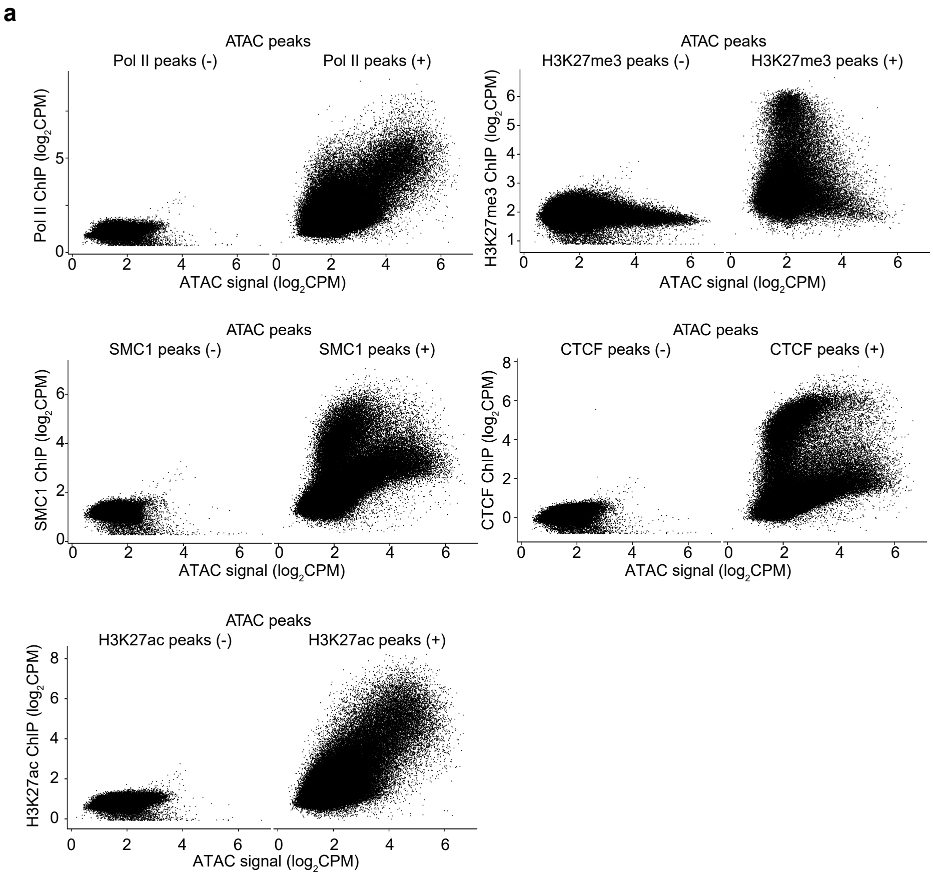


**Supplementary Fig. 2 Correlation between ATAC and ChIP signals. a.** Correlation between ATAC and indicated ChIP intensities at each ATAC peak with or without overlapping ChIP-seq peaks.

#### Supplementary Fig. 3

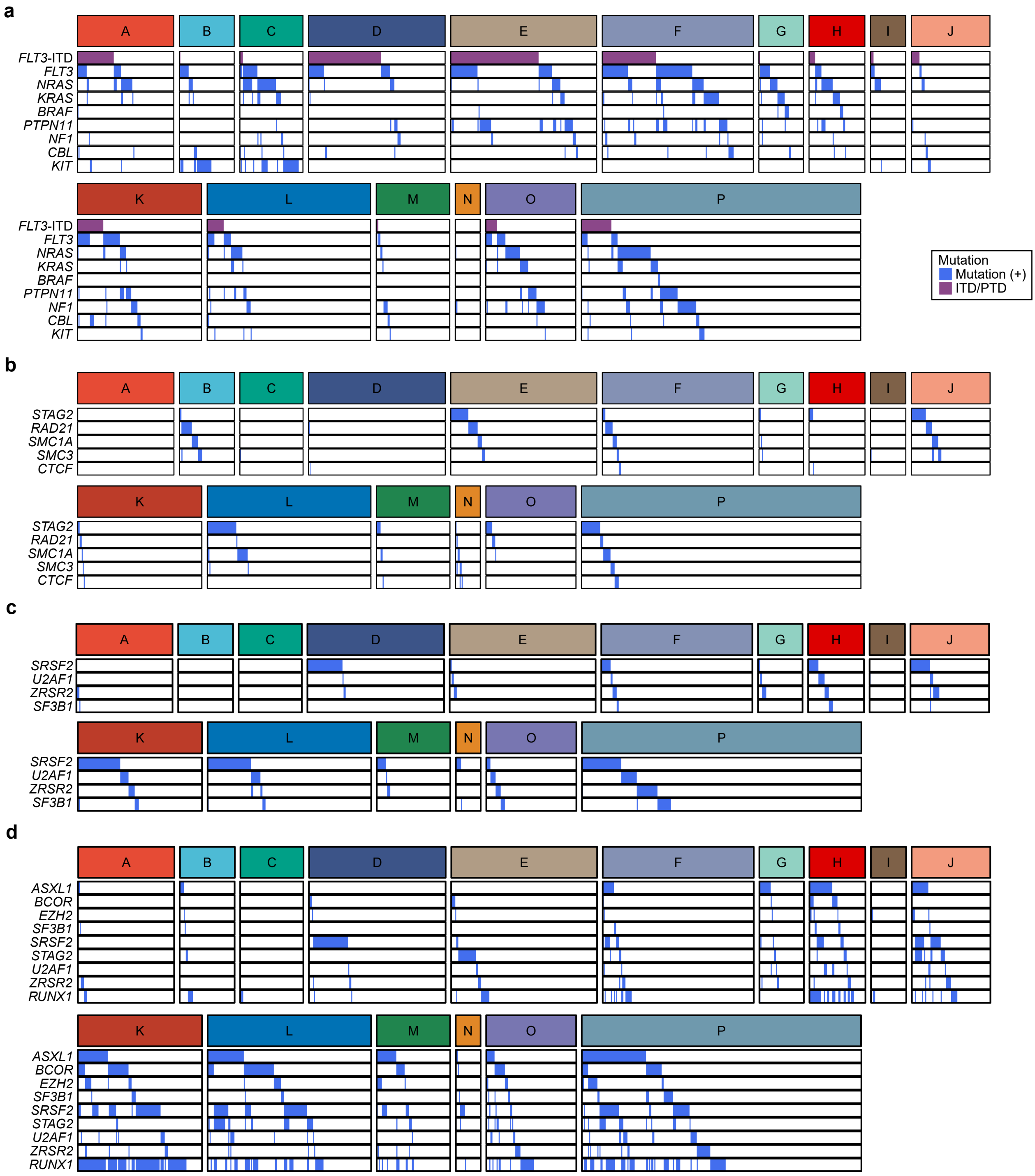


**Supplementary Fig. 3 Distribution of functionally categorized driver mutations across subgroups. a-d.** Summary of RAS pathway (**a**), cohesin (**b**), splicing factor (**c**), and MDS-related mutations (**d**) in each subgroup. Column represents patients, sorted by the presence of indicated genetic abnormalities.

#### Supplementary Fig. 4

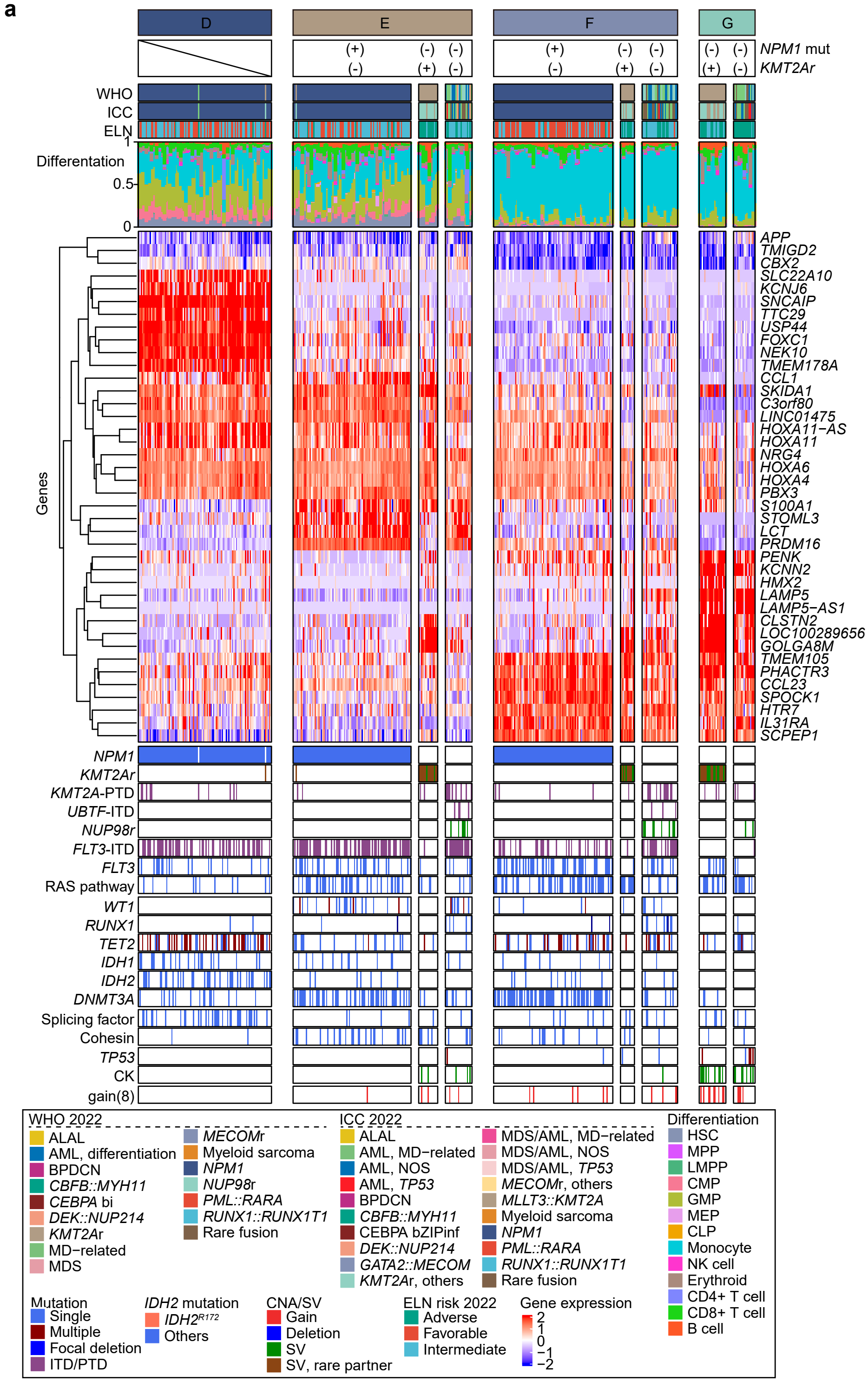


**Supplementary Fig. 4 Summary of HOX-related subgroups. a.** Summary of HOX-related subgroups according to driver mutations, such as *NPM1* mutations or *KMT2Ar*, with AML diagnoses, estimated differentiation status, normalized expression of representative genes, and genetic abnormalities. Each column represents one patient.

### Supplementary Data Fig. 5

**a**

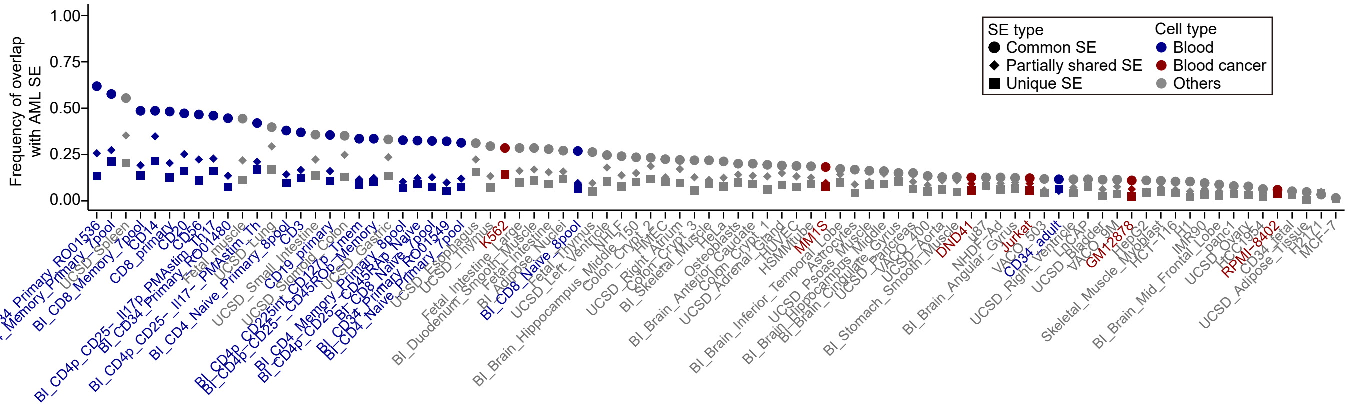


**Supplementary Fig. 5 Distribution of AML SEs across cell types and cancers. a.** Frequency of overlapped SEs from various cell types and cancers reported in a previous study[36] with common, partially shared, and unique SEs of AML (Fig. 2c).

#### Supplementary Data Fig. 6

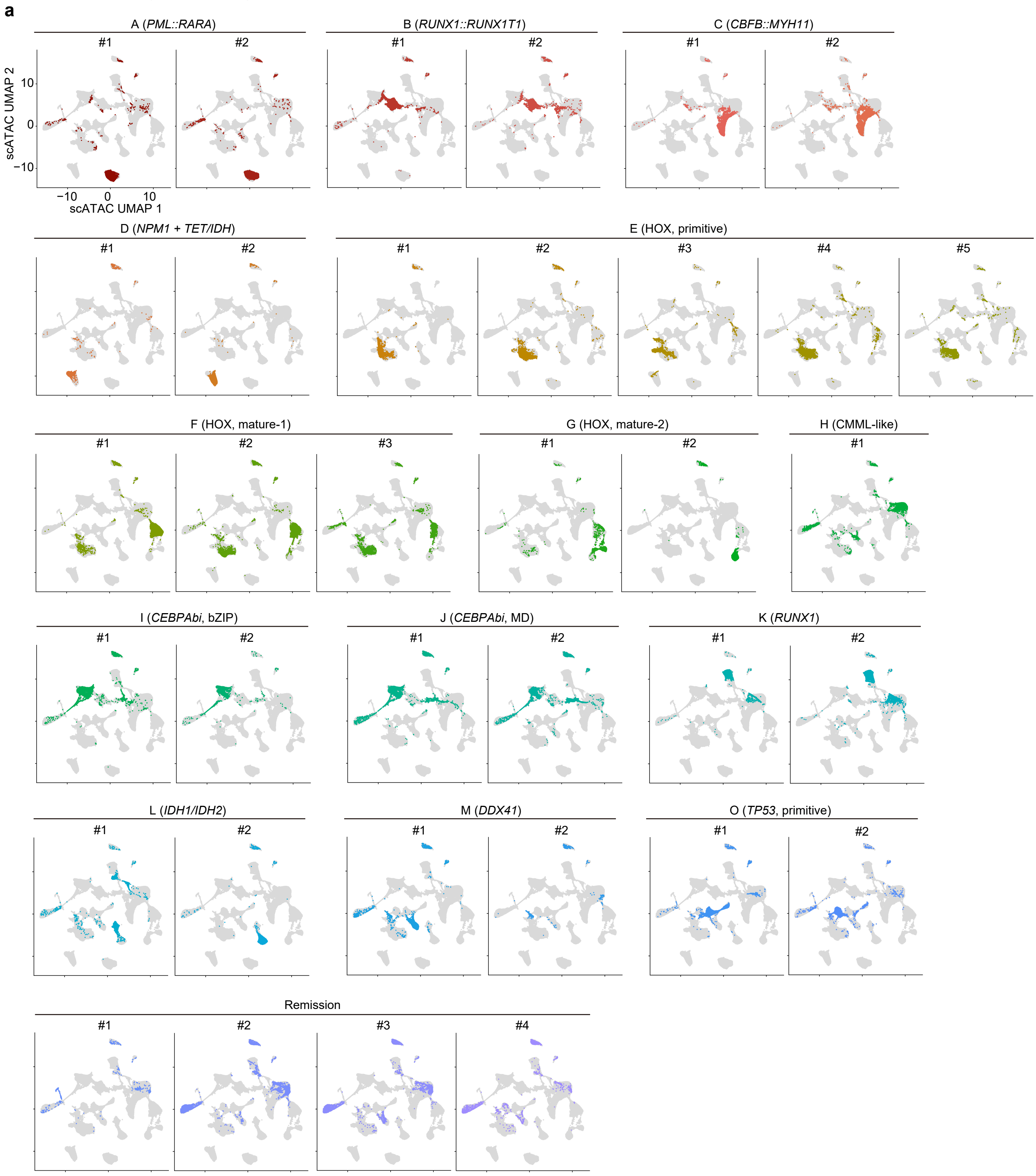


**Supplementary Fig. 6 scATAC-seq profiles for individual patient samples. a.** UMAP plots based on scATAC-seq profiles for each patient sample, indicated by color.

#### Supplementary Data Fig. 7

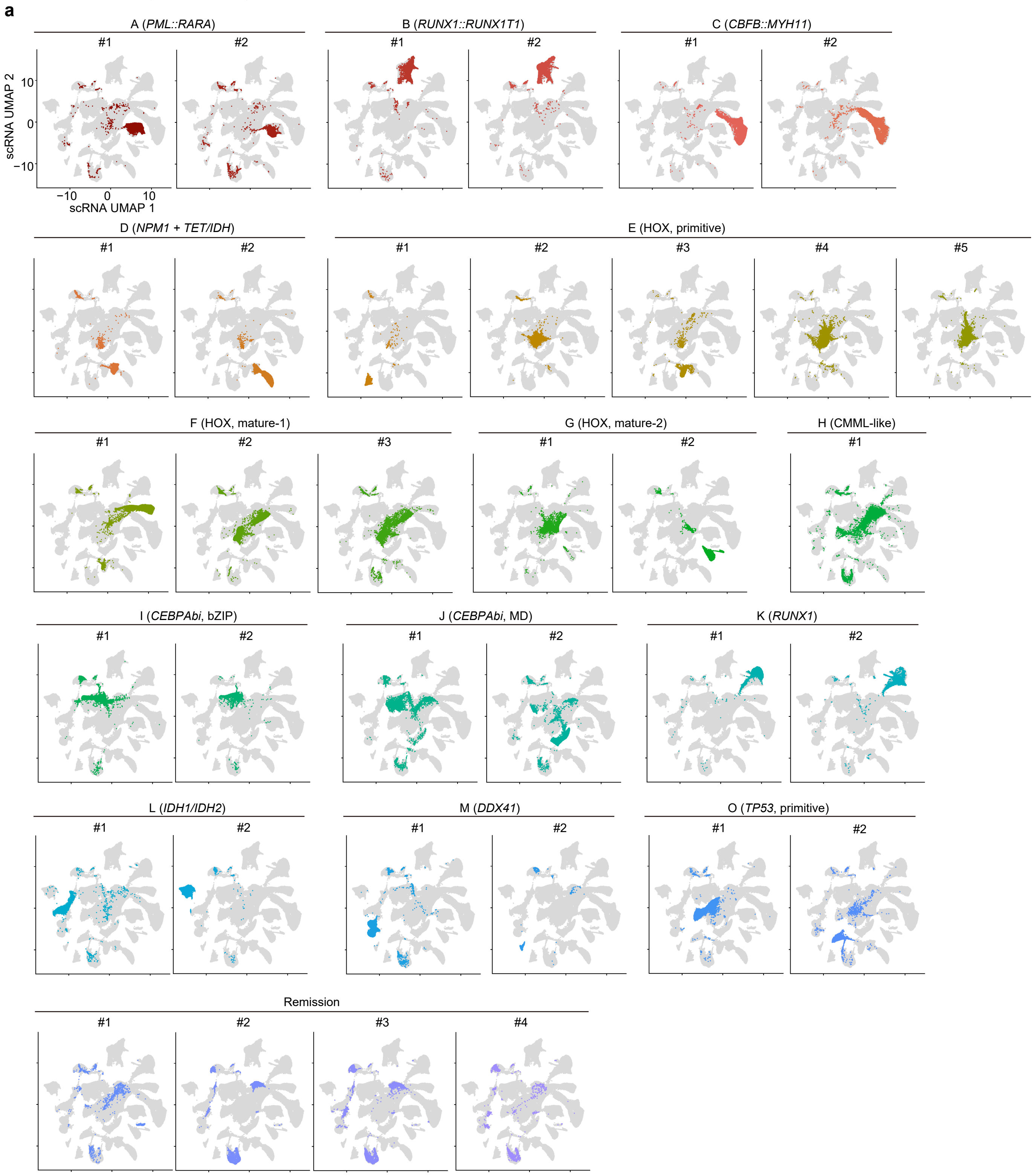


**Supplementary Fig. 7 scRNA-seq profiles for individual patient samples. a.** UMAP plots based on scRNA-seq profiles for each patient sample, indicated by color.

### Supplementary Data Fig. 8

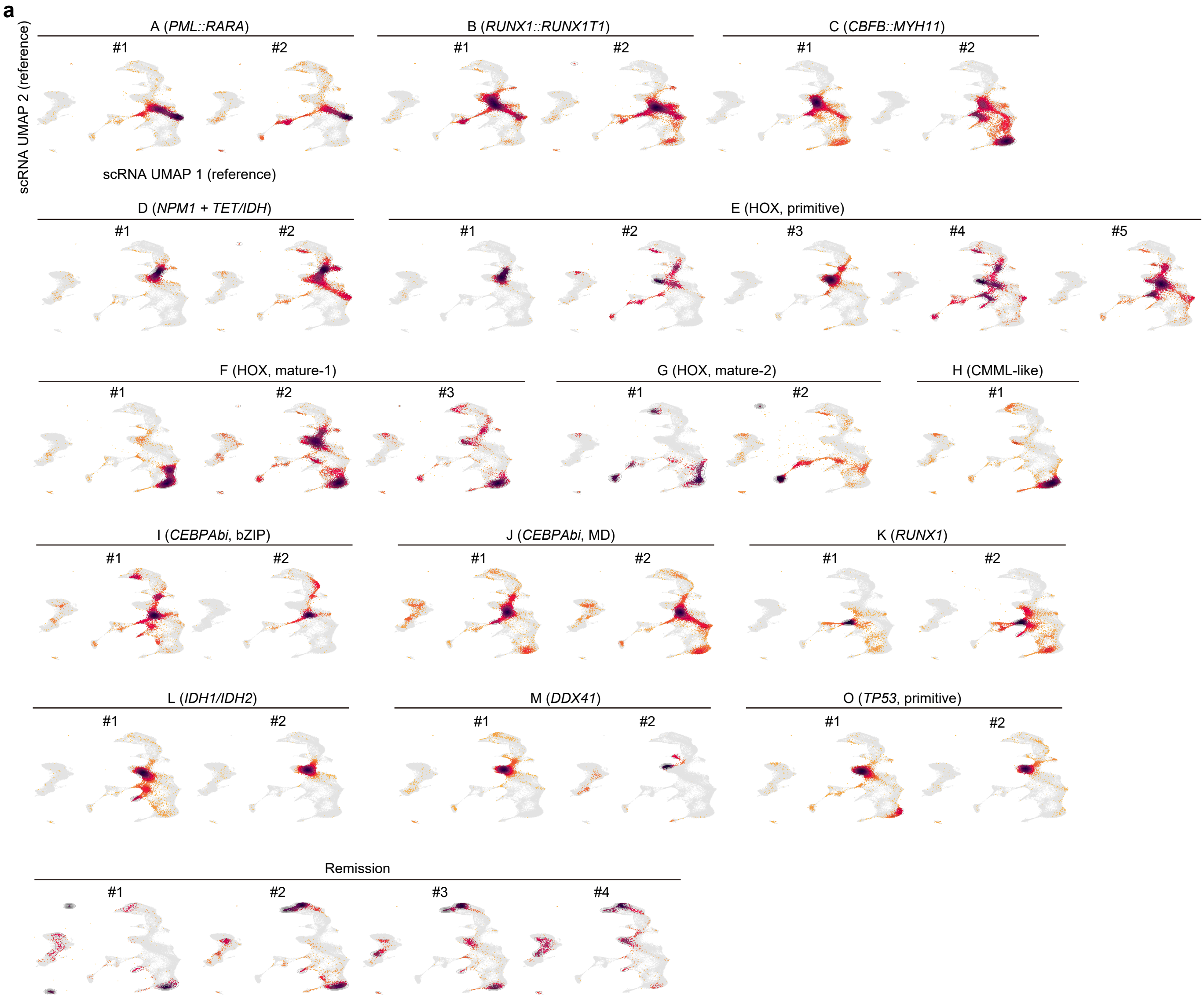


**Supplementary Fig. 8 Differentiation states of AML cells from individual patient samples. a.** UMAP plots of AML cells from each patient sample, indicated by color, mapped to the reference scRNA-seq UMAP.

### Supplementary Data Fig. 9

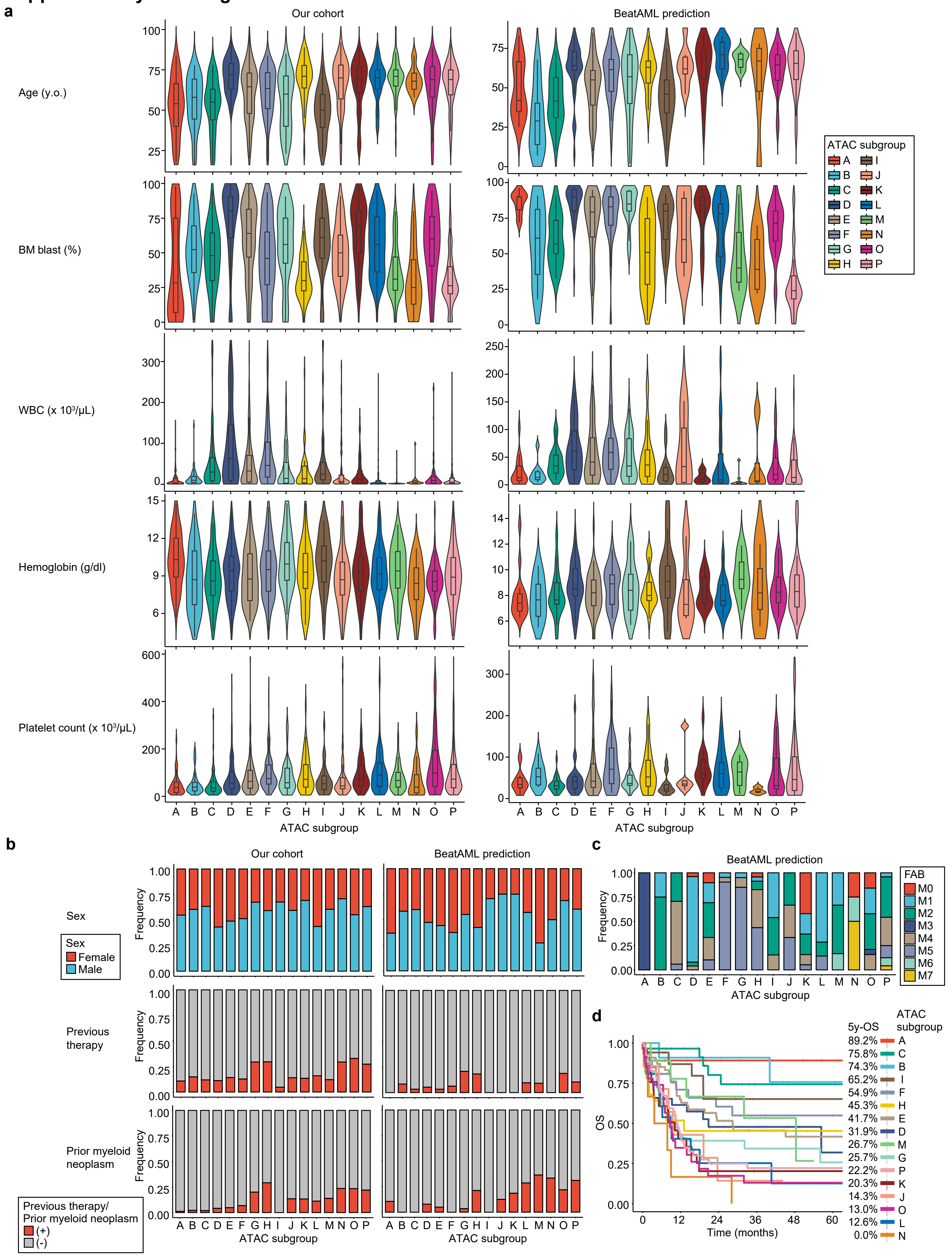

**Supplementary Fig. 9 Clinical features of ATAC subgroups.** **a-c.** Clinical parameters for ATAC subgroups in our dataset and predicted ATAC subgroups in the BeatAML dataset. **d.** Kaplan–Meier survival curves for OS of patients who received intensive chemotherapy in the BeatAML dataset, categorized by predicted ATAC subgroups. *P* value was calculated using the log-rank test.
